# Topsentinol L Trisulfate, a new Marine Natural Product, that Targets Basal-like and Claudin-low Breast Cancers

**DOI:** 10.1101/2020.04.17.047555

**Authors:** Nader N. El-Chaar, Thomas E. Smith, Gajendra Shrestha, Stephen R. Piccolo, Mary Kay Harper, Ryan M. Van Wagoner, Zhenyu Lu, Ashlee R. Venancio, Chris M. Ireland, Andrea H. Bild, Philip J. Moos

## Abstract

**Background:** Breast cancer is a heterogeneous disease. Genomic studies have revealed five different intrinsic subtypes - Luminal A, Luminal B, HER2-enriched, Claudin-low, and Basal-like. Patients diagnosed with Basal-like or Claudin-low breast cancer (BL-CL) suffer from poor prognosis and limited treatment options. Hence, there is an urgent need to identify a new therapeutic lead that can benefit patients with BL-CL breast cancer.

**Methods:** We used a step-wise screening approach to screen 2778 HP20 fractions from our Marine Invertebrate Compound Library (MICL) to identify compounds that specifically target BL-CL breast cancer. We also performed biochemical investigations to study the effect of the compound in the signaling pathway. Finally, we generated a drug response gene-expression signature and projected it against a human tumor panel of 12 different cancer types to identify other cancer types sensitive to the compound.

**Results:** We identified a previously unreported trisulfated sterol, topsentinol L trisulfate (TLT) that exhibits increased efficacy against BL-CL relative to Luminal/HER2+ breast cancer. Biochemical investigation of the effects of TLT on BL-CL revealed its ability to inhibit activation of AMPK and CHK1 and promote activation of p38. The importance of targeting AMPK and CHK1 in BL-CL was validated by treating a panel of breast cancer cell lines with known small molecule inhibitors of AMPK (Dorsomorphin) and CHK1 (Ly2603618) and recording the increased effectiveness against BL-CL compared to Luminal/HER2+ breast cancer. The TLT sensitivity gene-expression signature identified breast and bladder cancer as the most sensitive to TLT while glioblastoma multiforme as least sensitive.

**Conclusions:** Our study identified TLT, a previously uncharacterized trisulfated sterol, as a potential therapeutic selective against BL-CL. Our results also showed that inhibition of AMPK and/or CHK1 might be an effective therapy in BL-CL.

## INTRODUCTION

Gene-expression profiling has identified five molecular subtypes of breast cancer, known as Luminal A, Luminal B, HER2-enriched, Claudin-low, and Basal-like, with inter-subtype differences in incidence, survival and treatment response [1–5]. Among these, breast cancer patients diagnosed with the Claudin-low and Basal-like molecular subtypes exhibit particularly poor prognosis and suffer from limited treatment options [6]. Basal-like breast cancers represent 10-25% of all breast carcinomas, generally occurring at an early age (<40 years old), with higher frequency in women of African origin [7]. Approximately 50%-70% of all Basal-like cancers lack the expression of ER, PR, and HER2 and are therefore clinically described as being triple negative. Basal-like breast cancer manifests as a highly aggressive tumor that is responsive to chemotherapy [6]. However, patient prognosis remains poor with Basal-like cancers exhibiting a high recurrence rate and low patient survival [2]. At the molecular level, Basal-like breast cancers exhibit expression patterns as also observed in the basal epithelial layer of the skin and airways; this includes expression of high molecular weight cytokeratins 5, 6 and 17 and deficiencies in RB1, BRCA1 and TP53. Moreover, a high rate of aneuploidy is observed in these tumors, reflective of increased genetic instability [6, 7].

Interestingly, the Claudin-low group shares some similarities in gene-expression features with the Basal-like subtype such as low expression of HER2 and the luminal gene clusters, indicating genomic similarities between the two groups [8]. Moreover, like the Basal-like subtype, Claudin-low tumors are also triple-negative and have a poor prognostic outcome. However, Claudin-low breast cancers remain an individual group on their own, characterized by the minimal expression of several claudin genes, such as claudin 3, 4 and 7, which are involved in epithelial cell tight-tight junctions. These tumors also lack cell-cell junction proteins, such as E-cadherin, and almost always are characterized by having an intense immune cell infiltrate, stem cell properties, and features of epithelial-mesenchymal transition [3, 4, 7, 9, 10]. Due to the non-luminal molecular nature of these two subtypes, and the lack of known protein targets on these cancers, few effective treatment options are available. As such, there exists an urgent need to identify new therapeutic leads and potential targets that can improve patient prognosis.

The Marine Invertebrate Compound Library (MICL) is a unique resource that serves as a platform for discovery of novel small molecule-mediated biological activities in a variety of systems. MICL is derived from an extensive collection of small-molecule natural products isolated from over 1200 unique marine organisms (85% sponges from over 150 genera, 12% tunicates, 3% other phyla) collected from diverse locations around the world over the past twenty years [11, 12]. Natural products tend to be more complementary in shape to their targets [13] due to their development in a competitive ecological selection process that favors the production of compounds with strong biological activity [14–16]. Around 75% of all anti-cancer drugs developed between 1940 and 2014 were either derived from or inspired by natural products [17]. Several marine natural products, in particular, have been shown to exhibit anti-cancer properties, such as didemnin B, aplidine, and ecteinascidin-743, the latter of which succeeded in passing clinical trials in Europe and was approved by the European Commission for the treatment of refractory soft-tissue sarcomas in 2007 [18]. As of 2016, there were seven drugs approved by the United States Food and Drug Administrationthat are derived from marine natural products, four of which target cancers and 15 more marine natural products in clinical trials, of which 12 are anticancer [19]. In summary, murine natural products have proven their potential for development into clinically-useful drugs [18, 19].

In this study, we performed a step-wise screening approach with 2778 fractions from MICL and identified a previously uncharacterized trisulfated sterol, topsentinol L trisulfate (TLT), purified from a marine sponge *Topsentia* sp. (PNG07-3-073) collected from Papua New Guinea (Supplemental Figure 1). TLT, as well as its parent fraction and subfraction, exhibited increased effectiveness against BL-CL compared to Luminal and HER2+ subtypes. We showed that treating BL-CL cell lines with TLT leads to the inhibition of AMPK (Dorsomorphin) and CHK1 (Ly2603618), using biochemical and proteomic analyses. To validate the importance of inhibiting the activity of AMPK/CHK1 in BL-CL, we tested breast cancer cell lines with known small molecule inhibitors of AMPK and CHK1 and observed that they are significantly more effective against BL-CL than Luminal and HER2+ subtypes. Furthermore, overexpressing AMPK and/or CHK1 in BL-CL cell lines sensitized them to TLT treatment. We also generated a genomic gene-expression signature of TLT sensitivity and projected it against a panel of human patient tumors of twelve different cancer types, which identified breast and bladder cancer as the two cancers most sensitive to TLT. Overall, this study incorporates the genomic classification of breast cancer to high-throughput drug screening and identifies a novel small molecule, TLT, selective against BL-CL. Therefore, the work described here sheds light on the importance of targeting AMPK and/or CHK1 in this molecular subtype and suggests the potential of these proteins as therapeutic targets in BL-CL.

**Figure 1:**
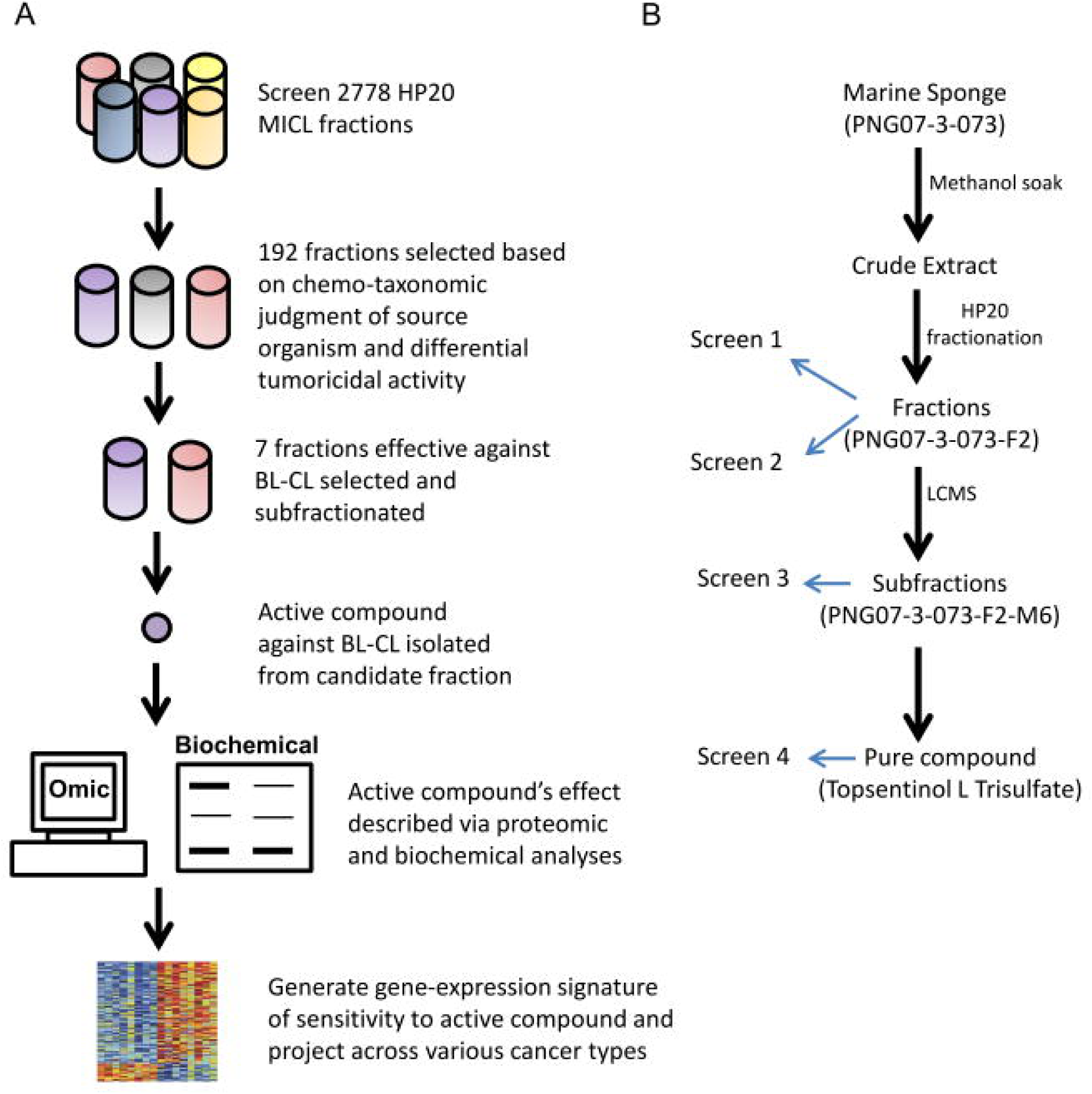
The overall design of the step-wise drug screen. **(A)** 2778 HP20 fractions were screened. One hundred ninety-two HP20 fractions were selected based on tumoricidal properties and chemo-taxonomic judgment of source organism. These fractions were then screened to identify BL-CL selective inhibitors. Seven candidate fractions were identified and further subfractionated. The subfractions were in turn screened for anti-BL-CL properties. After an anti-BL-CL subfraction candidate was identified, the active compound of the fraction was isolated, and its effect on BL-CL analyzed through proteomic and biochemical methods. A gene-expression signature of sensitivity to the active compound was then generated and used to project compound sensitivity across various cancer types **(B)** The path towards identifying TLT. Marine sponge PNG07-3-073 was diced and soaked in methanol to obtain a crude extract, which was fractionated on HP20SS resin. Among the five fractions obtained, only the F2 fraction exhibited tumoricidal activity (Screen 1). This activity was amplified against BL-CL (Screen 2). PNG07-3-073-F2 was then further fractionated, and the subfractions investigated for anti BL-CL activity, where the M6 fraction was identified as being BL-CL selective (Screen 3). Large-scale isolation of PNG07-3-073 ensued, culminating in the purification of TLT and its identification as the active compound in PNG07-3-073 responsible for anti-BL-CL effects (Screen 4).

## MATERIALS AND METHODS

### Cell Lines and Viability Measurement

Cell lines were obtained from ATCC and plated at 1500-2000 cells/well in 384-well plates in 5% FBS (Gibco/Life Technologies, Carlsbad, CA) growth media and 1x Anti-Anti (Gibco/Life Technologies, Carlsbad, CA). Cancer cell lines were cultured and maintained in a humidified environment at 37 °C and 5% CO2 in their respective media. The detailed description of the cell lines used for each screen is available in Supplementary Tables 1 and 2. Cells were treated for 72 hours, after which cell viability and growth were measured using CellTiter-Glo (Promega, Madison, WI). Cell viability scores were calculated by dividing the viability scores of the treatment by the control DMSO values.

### MICL Screens

For Screen 1, 2778 HP20 fractions of marine organisms from MICL were screened at a single dose (~1.5 µg/ml) against a panel of 16 (9 lung and 7 breast) cancer cell lines to determine their antitumor properties. We selected 107 fractions for further evaluation based on one or more of the following criteria: 1) all fractions with a standard deviation in viability of greater than 0.325; 2) lung selective fractions (25% or less viable cells in 3 or more lung cancer cell lines and 2 or fewer breast cancer cell lines); 3) breast selective fractions (25% or less viable cells after treatment in 3 or more breast cancer cell lines and 2 or fewer lung cancer cell lines); 4) generally active non-universally toxic fractions (25% or less viability in a minimum of 5 and maximum of 13 cell lines); 5) relatively less active fractions (40% or less viability in a minimum of 9 and maximum of 13 cell lines). For our second screen, we added an additional 85 HP20 fractions from MICL that were not included in the first screen but based on the chemo-taxonomic judgment of potential chemical similarity to the 107 fractions. These 192 HP20 fractions were then assayed at a single dose (~1.5 µg/ml) against a panel of 35 breast cancer and 37 lung cancer cell lines. This identified breast-selective fractions (fractions resulting in 25% or less cellular viability after treatment in 13 cell lines or more out of 35 breast cancer cell lines, and 12 cell lines or fewer lung cancer cell lines) and fractions effective against BL-CL (BL-CL vs Luminal/HER2+ breast cancer unpaired two-sample equal variance t-test < 0.05 with a positive average difference). Cell lines described as Basal-like or Claudin-low but being HER2+ were considered part of the Luminal/HER2+ group. Liquid-chromatography-mass-spectrum fractionation of the anti-BL-CL fractions following the MICL protocol [11, 12] resulted in 20 subfractions each that were assayed for effectiveness against BL-CL in a panel of 33 breast cancer cell lines. Once a candidate subfraction was determined based on the results of all three screens, large scale isolation and the purification of the active compound of that fraction was pursued (Supplementary Methods).

### Dose-Response Assays

Cell lines were plated as described above. TLT and halistanol sulfate were serially diluted 1:2 starting from 114.13 µM to the lowest dose of 3.57 µM and screened against a panel of 30 breast cancer cell lines. Dorsomorphin and Ly2603618 were serially diluted 1:3 starting from 90 µM to the lowest dose of 41.15 nM in RPMI media containing 5% FBS and 1x Anti-Anti and screened against a panel of 20 breast cancer cell lines, along with the combination treatment of Dorsomorphin and Ly2603618. For the combination treatment, an equal molar concentration of each compound was used. Cell viability was measured as described before. Doses were repeated in quadruplicates and averaged for a single value. EC50 values were calculated from dose-response curve data by plotting on GraphPad Prism 6.01 and using the equation: *Y* = 1/(1 + 10^((*logEC*50−*X*)∗*HillSlope*)^) with a variable slope (Ymin = 0 and Ymax = 1). Plots were forced to start from the x-axis by plotting for an x-intercept point.

### Reverse Phase Protein Array

Eight TLT-sensitive BL-CL cell lines (MDA-MB-157, MDA-MB-436, MDA-MB-468, MDA-MB-231, HCC38, HCC70, HCC1395, and HCC1143) were treated separately with TLT or DMSO at a concentration of 105 µM (approximate average EC75 across all 8 cell lines) for 6 hours, after which they were lysed according to the method detailed in supplementary methods. Reverse Phase Protein Array was performed at the University of Texas MD Anderson Cancer Center by the functional proteomics RPPA core facility according to their described methods and protocol [20, 21]. Two hundred seventeen different antibodies of phosphorylated and non-phosphorylated proteins were stained for and quantified (Supplementary Table 4).

### RNA Sequencing Data Acquisition

The same eight TLT-sensitive BL-CL cell lines were treated separately with TLT or DMSO for 6 hours, after which total RNA was extracted using the RNeasy Mini Kit (Qiagen, Venlo, Netherlands) with on-column digestion of the genomic DNA, as described in the manufacturer’s protocol. RNA sequencing was performed at the Huntsman Cancer Institute High Throughput Genomics Core Facility using 50-cycle, single-read sequencing (version 3) on an Illumina HiSeq instrument. To construct mRNA focused libraries from total RNA, the Illumina TruSeq RNA Sample Prep Kit (version 2) with oligo dT selection was used.

### TLT Sensitivity Signature Generation and Analysis

To process the mRNA sequencing data, we used the TCGA mRNA-seq Pipeline [22]. RNA sequencing reads for the treated and control samples were aligned using MapSplice v12_07 [23], quantified using RSEM [24], and gene counts were normalized using upper quantile normalization. This was the same methodology used to normalize the PANCAN12 TCGA dataset, which we obtained from TCGA fully processed for use in this analysis [22]. To generate a TLT sensitivity signature, we used the DESeq2 package (version 1.4.5) in the Bioconductor framework (version 2.14.0, version 3.1.0 of R) to identify genes that were significantly deregulated (adjusted p < 0.05) between the treated and control samples [25, 26]. One hundred forty-six genes were found to be significantly deregulated, out of which only 131 were found in the TCGA dataset. To use DEseq2, the reads had to be re-mapped using the Rsubread Bioconductor package. We used this package to map the reads to version hg19 of the human genome and to summarize the data to gene-level values [27]. We predicted TLT sensitivity for the PANCAN12 TCGA dataset [22] using the Bayesian binary regression algorithm version 2.0 (BinReg2.0) used as a MATLAB plug-in [28]. We used default parameters, except that our signature used 131 genes and one metagene. The probability output from the binary regression model was subtracted from one so that probabilities closer to one indicated a higher probability of sensitivity to the drug as previously described [29]. Prior to making the predictions, the data were log2 transformed and DWD normalized [30] to reduce biases that can result from differences in batch processing and platforms.

### Immunostaining

Four BL-CL cell lines sensitive to topsentinol L trisulfate (HCC70, HCC1143, MDA-MB-468, MDA-MB-436) were treated with 105 µM of TLT in 5% FBS RPMI media and 1x Anti-Anti for 6 hours. Proteins were extracted and western blots were run with the following primary antibodies – β-actin (#3700), β-tubulin (#2146), pRaptor-S792 (#2083), pCdc25c-S216 (#9528), pMAPKAPK-2-Thr334 (#3007), p38-T180/Y182 (#4511S), pChk1-S317 (#12302) and pAMPKα-T172 (#2535). All antibodies were obtained from Cell Signaling Technology (Beverly, MA).

### Statistical Analysis

To identify candidate fractions significantly more effective against BL-CL than Luminal/HER2+ breast cancer, preliminary statistical analysis was performed using the unpaired two-sample equal variance t-test built into the Microsoft Excel program. Final statistical assessment was performed for the fractions from the sponge PNG07-3-073 by re-analyzing statistical significance test based on the normality of the data. Gaussian distribution of the data was checked for using three different tests built into the GraphPad Prism 6 software: the D’Agostino-Pearson omnibus test, the Shapiro-Wilk test and the Kolmogorov-Smirnov test (with the Dallal-Wilkinson-Lilliefor corrected P value). Dot plots were then created using GraphPad Prism 6.01 and a standard two-tailed Mann-Whitney U-test was used to test for statistical significance, with the exception of the dot plot diagrams for halistanol sulfate, Dorsomorphin, Ly2603618, and Dorsomorphin + Ly2603618, where an unpaired t-test was used due to the normality of the data. To compare the difference of protein expression between treated cell lines and their DMSO controls and to test for the significance of the result, we used a two-tailed paired t-test.

## RESULTS

### Identification of topsentinol L trisulfate as a selective inhibitor of BL-CL

The goal of this study was to isolate a novel inhibitor of BL-CL, describe the pathways effectively blocked by the compound, and project treatment efficacy across other cancer types. We aimed to achieve this through a stepwise approach by 1) screening MICL for fractions that exhibited tumoricidal properties; 2) selecting candidate fractions that displayed effectiveness against BL-CL; 3) separating the active compounds in the fractions; 4) identifying the active compound with anti-BL-CL properties; 5) describing the cell signaling effects of the compound on BL-CL; 6) projecting TLT sensitivity across human tumors of various cancer types using a gene-expression signature (Figure 1A).

We used a bioassay-guided fractionation approach to identify potential compounds with tumoricidal activity, beginning by screening 2778 HP20 fractions from MICL against a panel of 16 cell lines (Figure 1B “Screen 1”). These fractions represent complex mixtures and served as a starting point to identify promising hits [11, 12]. We selected 107 fractions based on differential tumoricidal activity, eliminating all fractions that were universally toxic or showed minimal anti-cancer activity. We added 85 previously-unscreened HP20 fractions from MICL based on chemical similarity to the 107 selected fractions guided by chemo-taxonomic considerations of source organisms [11, 12]. The combined 192 fractions were screened against a panel of 35 breast cancer and 37 lung cell lines to identify fractions more effective against breast cancer, and in particular against BL-CL (Screen 2). Of these, 34 fractions were selective for breast cancer and seven of those displayed subtype selectivity against BL-CL (Figure 1B “Screen 2”). These seven fractions were further fractionated by LCMS into 20 subfractions (M1-M20) and screened at two doses against a panel of 33 breast cancer cell lines (Figure 1B “Screen 3”). This screen identified five subfractions with significant selectivity against BL-CL, out of which three subfractions originated from the marine sponge PNG07-3-073 F2 fraction. We analyzed the results of all three screens retrospectively and observed that PNG07-3-073–F2 was initially identified as a candidate with increased tumoricidal activity against BL-CL (Figure 2A, Figure 1B). The HP20 fractionation of the sponge also resulted in 4 other fractions, FW, F1, F3, and F4; however, none exhibited sufficient tumoricidal activity and therefore did not proceed past the first screen. The F2 was the only fraction from the *Topsentia* sponge PNG07-3-073 to exhibit anti-BL-CL activity (Figure 1B). This activity was maintained after further fractionation of F2 via LCMS into 20 subfractions. Out of 20 subfractions, the M6 subfraction, in particular, displayed significant effectiveness against BL-CL (Figure 2B, Figure 1B).

**Figure 2:**
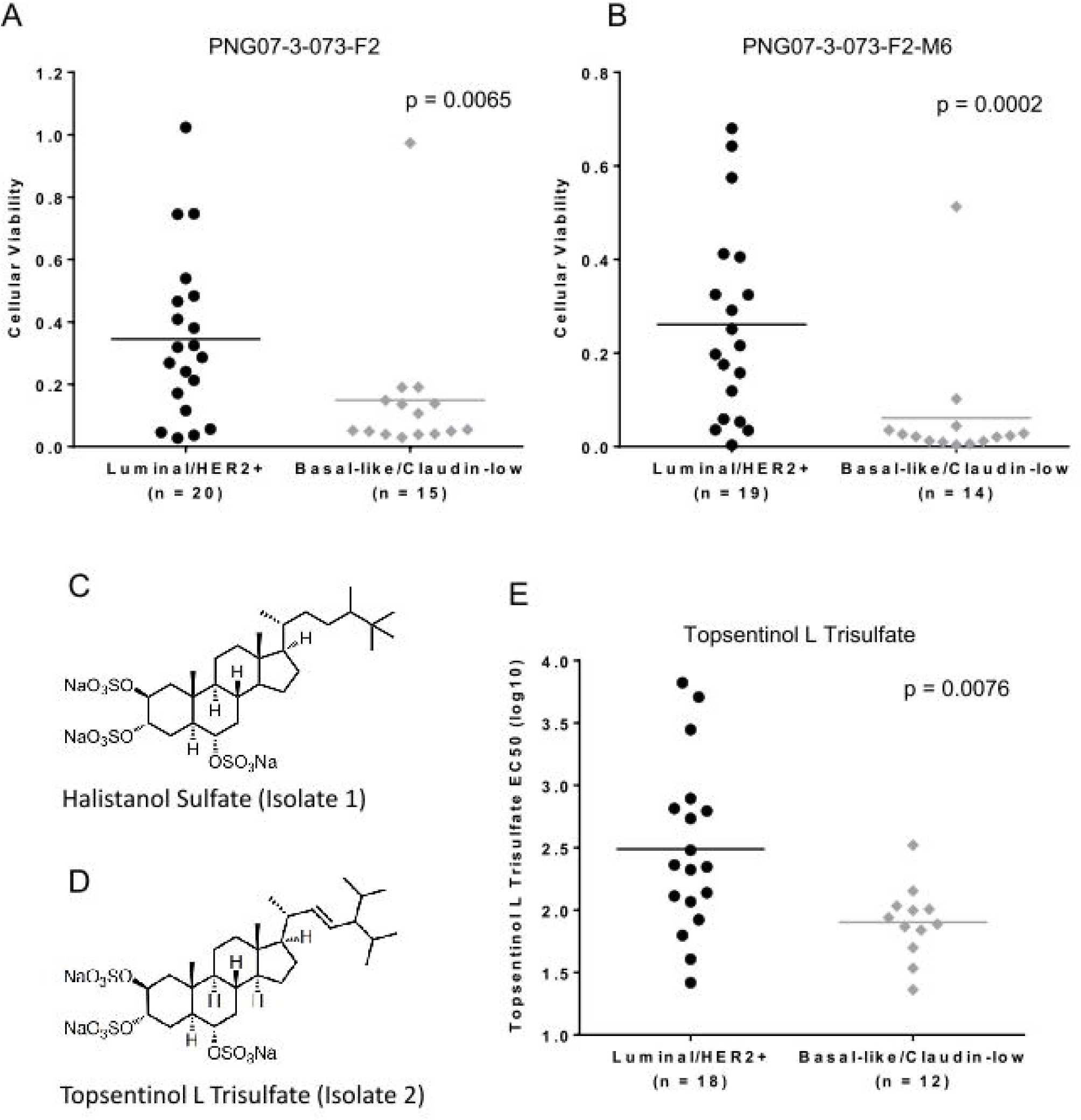
Topsentinol L trisulfate, isolated from the marine sponge *Topsentia* sp. (PNG07-3-073), is selective against BL-CL. Cell lines were treated with TLT and viability was measured. Every dot represents a cell line, with the y value representing the cell line’s viability or compound EC50 post treatment. The horizontal line indicates the mean for every group. BL-CL cell lines exhibit significantly lower cell viability when treated with **(A)** PNG07-3-073-F2 and **(B)** PNG07-3-073-F2-M6 than Luminal/HER2+ cell lines. **(C)** Chemical structure of halistanol sulfate **(D)** Chemical structure of topsentinol L trisulfate **(E)** Response to topsentinol L trisulfate treatment as measured by compound EC50. BL-CL cell lines are significantly more sensitive to TLT than Luminal/HER2+ cell lines.

Our next step was to proceed with the identification of the active compounds in the M6 subfraction with the aim of isolating any compounds inducing this response (Figure 1A). We performed a scaled-up extraction of bulk PNG07-3-073 sponge and purified two compounds: isolate 1 and 2. 1D NMR analysis (Supplementary Table 3) identified isolate 1 as the previously reported metabolite halistanol sulfate [31] (Figure 2C, Supplementary Figure 2A-B) and was validated using low-resolution mass spectroscopy (Supplementary Figure 3A). Isolate 2 was identified via 1D (Supplementary Figure 2C-2D) and 2D (Supplementary Figure 4) NMR analysis (Supplementary Table 3) as a previously uncharacterized sulfated sterol identical to the known compound topsentinol L but with three sulfate groups [32] (Figure 2D). The structure of the compound was corroborated by the data obtained through low-resolution mass spectroscopy (Supplementary Figure 3B). We named this compound topsentinol L trisulfate, or TLT (Figure 2D). Both structures were validated by high-resolution electrospray ionization mass spectrometry (Supplementary Methods). Halistanol sulfate and TLT were then screened against a panel of 30 breast cancer cell lines and tested for effectiveness against BL-CL. TLT showed significant tumoricidal activity against BL-CL breast cancer compared to other subtypes (p = 0.0076, Figure 2E, Figure 1B). Halistanol sulfate did not exhibit such activity (p = 0.247, Supplementary Figure 5). Through a multi-step screening and fractionation process, we identified a novel sulfated sterol, TLT, which exhibits significant subtype selectivity against BL-CL.

**Figure 3:**
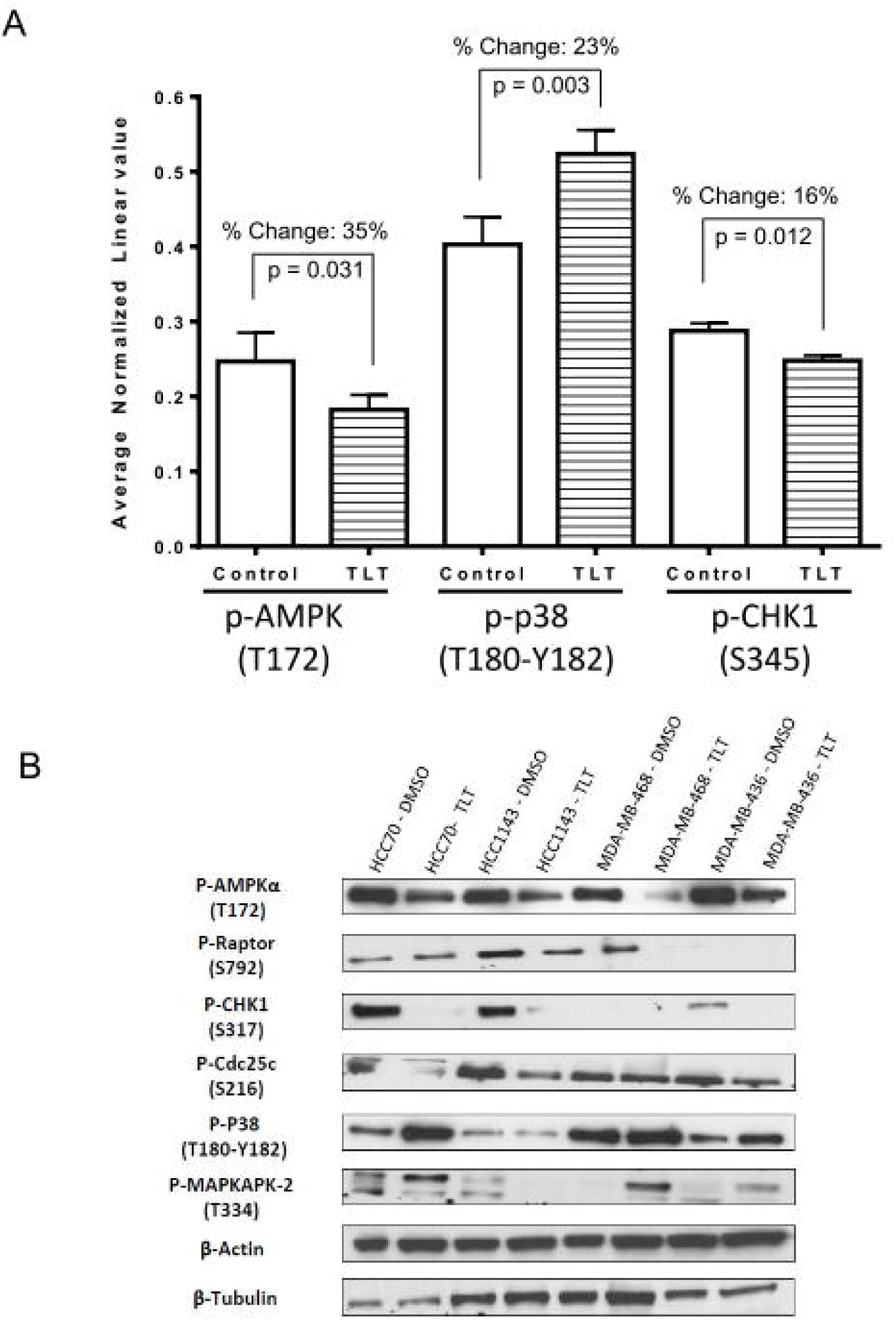
TLT inhibits AMPK and CHK1 but activates p38. **(A)** A panel of 8 BL-CL cell lines were treated with TLT at a 105 µM dose (approximate average EC75 dose across all 8 lines) for 6 hours and compared to DMSO control via RPPA that investigated 217 proteins. Eight proteins displayed 15% or more significant deregulation in protein levels, among which, AMPK, CHK1 and p38 are displayed here in bar graphs. Error bars represent SEM. **(B)** Observations made for p38, AMPK and CHK1 in the RPPA experiment were validated by Western Blotting. A panel of four BL-CL cell lines were treated similarly with TLT at a 105 µM dose for 6 hours and compared to DMSO control. TLT treatment inhibits AMPK and CHK1, and activates p38. β-actin and β-tubulin were used here as protein loading controls.

**Figure 4:**
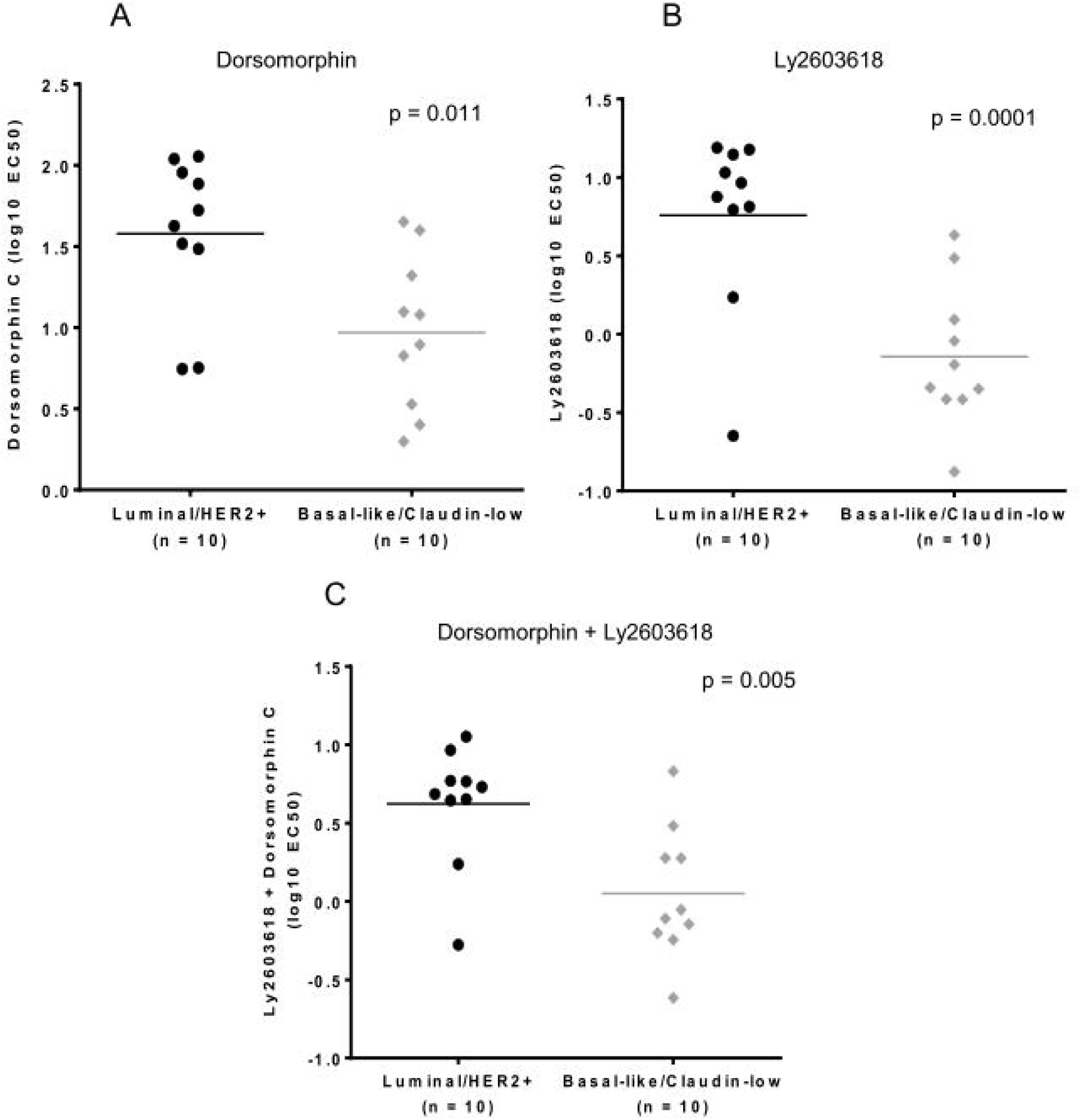
Single or dual inhibition of AMPK and CHK1 is selective against BL-CL. Cell lines were treated and scatter dot diagrams of treatment EC50 values were plotted. Every dot represents a cell line, with the y value representing the cell line’s treatment EC50 and the horizontal line indicating the mean for each group. Cells lines were treated with the specified inhibitor for 72 hours. BL-CL cell lines are significantly more sensitive to **(A)** AMPK inhibition through Dorsomorphin **(B)** CHK1 inhibition through Ly2603618 and **(C)** the combined inhibition of both AMPK and CHK1 through concurrent treatment with Dorsomorphin and Ly2603618.

**Figure 5:**
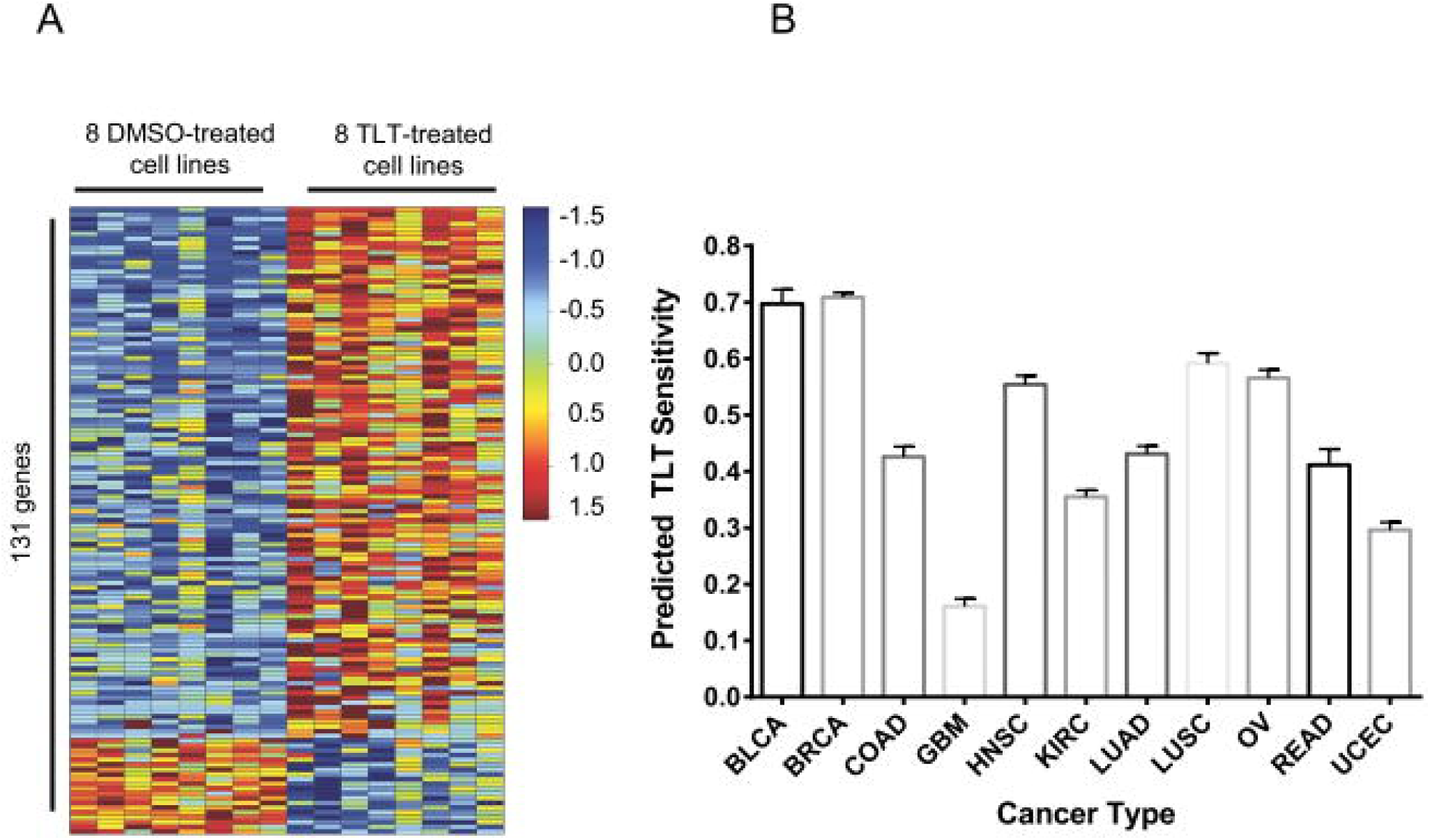
TLT sensitivity is predicted in breast and bladder cancer using gene-expression signature analysis. **(A)** The gene-expression signature for TLT sensitivity was generated by treating eight BL-CL cell lines with either TLT or DMSO. The heatmap columns are the eight cell lines making up eight controls on the left (treated with DMSO) and eight treated samples on the right (treated with TLT). Each row represents a gene that is part of the signature. There are a total of 131 genes making up the signature. Red indicates upregulation while blue indicates downregulation of the gene. **(B)** The TLT sensitivity signature was used to project the sensitivity of twelve different cancers. Results are shown in a bar graph where the x-axis represents the twelve cancer types assayed and the y-axis represents the predicted score of TLT sensitivity (minimum = 0, maximum = 1). Each column portrays the mean of the TLT sensitivity scores across the samples in a particular cancer type. The error bars indicate SEM. Breast and bladder cancer were predicted to be the most sensitive to TLT treatment, with glioblastoma being the least sensitive. Legend = BLCA: bladder urothelial carcinoma. BRCA: breast invasive carcinoma. COAD: colon adenocarcinoma. GBM: glioblastoma multiforme. HNSC: head and neck squamous cell carcinoma. KIRC: kidney renal clear cell carcinoma. LUAD: lung adenocarcinoma. LUSC: lung squamous cell carcinoma. OV: ovarian serous cystadenocarcinoma. READ: rectum adenocarcinoma. UCEC: uterine corpus endometrial carcinoma.

### Topsentinol L trisulfate treatment inhibits AMPKα and CHK1 but activates p38

Our next goal was to analyze and describe the signaling effects that are induced by TLT in cancer cells. For this purpose, we treated eight TLT-sensitive BL-CL cell lines with TLT or DMSO control and screened for 217 phosphorylated and non-phosphorylated protein changes using reverse-phase protein array (RPPA) [20]. From this screen, we identified 21 proteins that exhibited a 15% or more upregulation or downregulation in protein level, with only eight of these proteins exhibiting statistically significant changes (Supplementary Table 4). To focus our research and narrow down our investigation, we selected three proteins, AMPK, CHK1, and p38 based on fold change and statistical significance. AMPK phosphorylation recorded the largest statistically significant change compared to DMSO control of all proteins, with 35% reduction in Thr172 phosphorylation, which is required for AMPK activation [33] (p = 0.031, Figure 3A, Supplementary Table 4). Validating AMPK inhibition, phosphorylation of ACC, a direct downstream effector of AMPK [34], was also significantly reduced, recording a 23.6% reduction in Ser79, supporting the observation of AMPK inhibition (Supplementary Table 4, p = 0.024). We also observed significant changes in phosphorylation of CHK1, recording a 16% downregulation in the nuclear localization mark Ser345 [35] (p = 0.012, Figure 3A, Supplementary Table 4). CHK1 has recently been suggested to represent a therapeutic target in triple-negative breast cancer (TNBC) [36], therefore making its inhibition following TLT treatment a potential cause of treatment effectiveness. Upregulation of p38 phosphorylation was the most significant change observed upon treatment with TLT, with a 23% increase in Thr180-Tyr182 phosphorylation (p = 0.003, Figure 3A, Supplementary Table 4). The activation of p38 has been shown to lead to the induction of apoptosis [37].

In order to validate the observed deregulation of AMPK, p38, and CHK1 following TLT treatment in the RPPA screen, we analyzed phosphorylation levels by Western Blotting. The most consistent downregulation observed was inhibition of AMPK activation across all four cell lines (Figure 3B). CHK1 activity measured by the phosphorylation of the activation site of the protein, Ser317, was inhibited by TLT [38] (Figure 3B). In addition, the phosphorylation of p38 was also upregulated in three out of four cell lines, confirming previous results (Figure 3A-3B). Validating the inhibition of these three kinases, Raptor, Cdc25c and MAPKAPK-2, substrates of AMPK, CHK1 and p38 respectively [36, 37, 39], were observed to be inhibited in at least 2 of the cell lines tested following TLT treatment (Figure 3B). These data describe the landscape of proteomic changes induced following TLT treatment in BL-CL cells.

### Inhibition of AMPK and CHK1, alone or in combination, is effective against BL-CL

Due to the consistency with which AMPK (either the protein itself or its downstream effector ACC) and CHK1 (the inhibition of two separate activation marks Ser345 and Ser317) are inhibited, as well as the recent discovery of CHK1 treatment efficacy against TNBC [36], we decided to test the relevance of the inhibiting AMPK and CHK1 in BL-CL. We investigated the effects of AMPK, CHK1 and AMPK1+CHK1 small molecule targeted inhibition on a panel of twenty breast cancer cell lines. We screened the panel with Dorsomorphin, an AMPK inhibitor [40], Ly2603618, a CHK1 inhibitor in phase two clinical trials [41], and the combination of both. Our results showed that either treatment strategy is significantly more effective against the BL-CL subtype than Luminal/HER2+ breast cancer (Figure 4). Dorsomorphin was more than four times more effective against BL-CL (average EC50 = 9.33 µM) compared to Luminal/HER2+ breast cancer (average EC50 = 37.87 µM), which was a significant difference (p = 0.011, Figure 4A). Ly2603618, on the other hand, was almost eight times more toxic against BL-CL (average EC50 = 0.72 µM) in comparison to Luminal/HER2 breast cancer (average EC50 = 5.73 µM), which was also a significant difference (p = 0.001, Figure 4B). Interestingly, the combination therapy was also significantly more effective against BL-CL (p= 0.005) (Figure 4C). To confirm that AMPK and CHK1 are part of the killing mechanism, we overexpressed AMPK and/or CHK1 in two BL-CL cell lines and performed dose-response assays with TLT to study the effect of AMPK and CHK1 abundance on drug response. We observed that the overexpression of AMPK, CHK1 or both concurrently led to the sensitization of the cell lines to TLT treatment (Supplementary Figure 6). Indeed, the increase in the availability of AMPK and/or CHK1 lowered the EC50 of TLT across both cell lines (Supplementary Table 5), further linking the TLT killing mechanism to the AMPK and CHK1 proteins. These results validate the importance of the inhibitory effects of TLT on AMPK and CHK1 in BL-CL and suggest the potential use of AMPK or CHK1 inhibition as a treatment.

### TLT sensitivity signature predicts breast and bladder cancer response in human tumors

Next, we sought to identify classes of solid tumors that are most sensitive to TLT using an unbiased computational approach. The gene-expression profiles of cancer cells are a valuable tool in the comprehension of transcriptional changes indicative of treatment. These profiles enable the identification of drug sensitivity across various cancer types. Such expression profiles may be used to characterize the genes whose expression is indicative of drug response [29]. Accordingly, we generated a TLT sensitivity signature that reflected the genomic changes in eight TLT-sensitive BL-CL cell lines (Figure 5A). We treated the cell lines with either TLT or DMSO control and used RNA-sequencing to profile the samples. We identified 131 genes that were significantly upregulated or downregulated and incorporated the expression values for these genes into a predictive signature (Figure 5A). We then used this signature to predict drug sensitivity for the tumors from the PANCAN12 gene-expression dataset, which contains expression profiles for 12 cancer types [22, 42]. The outcome of this process is a probability for each tumor sample, indicating how likely each tumor would respond to TLT treatment. Breast (mean = 0.71) and bladder cancer (mean = 0.69) were predicted to be most sensitive to TLT (1.00 is highest possible sensitivity, and 0.00 is lowest), while glioblastoma was predicted to be least sensitive (mean = 0.16, Figure 5B). These results are in line with the *in vitro* observations of TLT effectiveness against breast cancer (BL-CL in particular) and suggest future investigation of TLT as an effective therapeutic lead against bladder cancer.

## DISCUSSION

We have identified and described a previously uncharacterized trisulfated sterol that we have named topsentinol L trisulfate (TLT) that exhibits increased tumoricidal activity against BL-CL. Interestingly, halistanol sulfate, another trisulfated sterol isolated from the same marine organism as TLT, did not exhibit similar activity against BL-CL (Supplementary Figure 5). This could potentially be attributed to the differences in the side chains of these two compounds, which are otherwise structurally identical (Figure 2C, 2D).

Furthermore, we describe the treatment effect of TLT on BL-CL, highlighting the particular changes in the activation of AMPK, CHK1, and p38. AMPK is a heterotrimeric serine/threonine kinase complex that is regulated by adenylate levels in the cell and functions as part of an evolutionarily conserved energy-sensing pathway [43, 44]. The effective result of AMPK activation is the avoidance of bioenergetic catastrophy and cell death through the conservation of cellular energy [44]. Interestingly, the role of AMPK in cancer is complex, as AMPK can exert pro- or anti-tumor effects based on cell context. AMPK is central to a tumor suppressor network, the LKB1-AMPK-TSC-mTOR signaling cascade, known to regulate cell growth and proliferation in response to stress [45]. Conversely, retaining continuous activation of AMPK leading to an enhanced ability to adapt to metabolic stress may function to promote tumor survival and growth. For example, the activation of AMPK in response to stresses such as hypoxia and nutrient deprivation provides cancer cells with the metabolic flexibility needed for survival [44]. These dueling roles of AMPK highlight the complexity and dichotomy of the kinase’s role in cancer cells. AMPK agonists acting as anti-cancer agents have been suggested through the use of the therapeutic biguanides, metformin, and phenformin. Metformin is currently used to treat Type II diabetes and has been associated with a significantly lower cancer incidence in patients relative to those using other medications to manage their diabetes [40, 46]. However, recent work has indicated that the anti-tumorigenic effects of metformin and another known AMPK agonist, AICAR, are due to AMPK-independent effects [39]. Interestingly, other studies have implicated AMPK as a mediator of cellular proliferation and survival, showing the promising effect of AMPK inhibition as cancer therapy [47, 48]. We observe similar effects against breast cancer, and particularly the BL-CL subtype that exhibits higher sensitivity against Dorsomorphin than Luminal/HER2+ breast cancer (Figure 4A). This observation is in line with the AMPK inhibitory effects induced in BL-CL breast cancer when treated with TLT (Figure 3A-3B).

Another aspect of the inhibitory effects promoted by TLT treatment was the downregulation of CHK1 activation. Upon cellular exposure to various genotoxic stresses, CHK1 is activated by ATR-mediated phosphorylation following DNA-damage leading to the phosphorylation of cdc25. CHK1 assumes the role of the major cell-cycle checkpoint kinase mediating S- and G2-arrest [36]. In BL-CL, the rationale of CHK1 targeted therapy is supported by the documented evidence of alterations in the DNA damage repair machinery through either the high rate of BRCA or p53 mutations [6, 7, 9]. Therefore, another loss of a DNA damage repair component may lead to the cell’s inability to properly fix chromosomal damage and enter apoptosis. Indeed, Albiges, *et al.* have shown that CHK1 is a potential therapeutic target in TNBC, with CHK1 inhibition observed to induce mitotic cell death in TNBC cell lines [36]. Our observation of increased sensitivity of BL-CL to CHK1 inhibition is concordant with the TNBC subtype (Figure 4B).

Similarly, we have shown that treating BL-CL breast cancers with TLT leads to the significant activation of p38 (Figure 3A, Supplementary Table 4). p38 plays the role of a signal transduction mediator and is linked to inflammation, cell cycle, cell death, cell differentiation, senescence and tumorigenesis (37). A number of studies have been shown that the activation of p38 leads to apoptosis in various cells (49, 50).

The inhibitory effects of TLT on AMPK and CHK1 shed light on the potential therapeutic benefit of AMPK and/or CHK1 inhibition on BL-CL. However, additional work is required to validate these findings. Although Dorsomorphin is a potent AMPK inhibitor, studies have shown this compound to exhibit high affinity towards other proteins such as BMP and inhibit several other kinases at a Km lower than AMPK [49], prompting us to validate the role of TLT-mediated inhibition of AMPK via genetic means (Supplemental Figure 6). Furthermore, TLT also strongly inhibited AKT, MAPK, RAF, FAK, STAT3, p70S6K, FASN and PDK1 (Supplementary Table 4), of which all of these are likely to have strong inhibitory effects on cell viability, and therefore cannot be discounted to participate in the killing mechanism alongside AMPK and CHK1 inhibition. Our work highlights the effects of TLT on BL-CL and implicates a number of proteins involved in its downstream repercussions but does not address the specific mechanism of action leading to these effects.

## CONCLUSIONS

Our work describes the projected efficacy of TLT against a variety of human tumors, highlighting the optimal effect of the compound against breast and bladder cancer. In a TCGA study characterizing the molecular landscape of urothelial bladder carcinoma, a p53 mutation rate of 49% was recorded in the samples tested [50], drawing a similarity to the common observation of p53 mutation in BL-CL. Remarkably still, among the key pathway nodes deregulated in bladder cancer, the LKB1/STK11-TSC-mTOR node was among the most commonly deregulated. LKB1, the activator of AMPK, was recorded to contain copy number alternations (CNAs) in 11% of all cases. TSC1 and TSC2 recorded CNAs in 16% and 9% of all cases, as well as inactivating mutations in 8% and 2% respectively [50]. Thus, one hypothesis is that the inhibitory effect of TLT on CHK1 and AMPK (and subsequently the LKB1-AMPK-TSC-mTOR node) could lead to an equally effective response against bladder cancer. Further work is needed to elucidate the exact mechanism of action of TLT and its projected effectiveness against bladder cancer.

In this study, we have described the discovery and identification of a previously unreported sulfated sterol, as well as its signaling effect on BL-CL. We have described two potential therapeutic targets of BL-CL that can be exploited for the benefit of treatment efficacy. This lays the groundwork necessary for the exploration of AMPK and CHK1 as potential targets of BL-CL treatment, with a need to further characterize and delineate the role of TLT as an investigational anti-BL-CL compound.

## List of abbreviations

AMPK: AMP-activated protein kinase
BL: Basal-like
BRCA1: BRCA1, DNA repair associated
CL: Claudin-low
CHK1: checkpoint kinase 1
EC: Effective concentration (generally followed by a number for the percent activity)
ER: estrogen receptor
HER2: human epidermal growth factor receptor 2
HP20: synthetic polymer resin for high-performance liquid chromatography
LKB1/STK11: liver kinase B1, serine/threonine kinase 11
MICL: Marine Invertebrate Compound Library
PR: progesterone receptor
RB1: retinoblastoma transcriptional corepressor 1
TLT: topsentinol L trisulfate
TP53: tumor protein p53

## DECLARATIONS

### Ethics approval and consent to participate

Not applicable.

### Consent for publication

Not applicable.

### Availability of Data

The raw and processed data are available in the Gene Expression Omnibus under the series accession number GSE142833.

### Authors contributions

PJM, AHB, and CMI contributed to the study design. CMI, MKH, and RMV established and maintained the MICL, TES and MKH purified halistanol sulfate and TLT, TES and ZL performed structure elucidations, and TES prepared supplemental material pertaining to chemical structures. PJM, NNE-C and GS performed to the drug screen and NNE-C and GS contributed to the cell biology experiments. PJM, AHB, NNE-C and SRP contributed to the data analysis. SRP was responsible for the bioinformatics analysis. NNE-C drafted the manuscript and all authors contributed to the editing of the manuscript.

## Acknowledgments

This study was supported by the National Institutes of Health (R01GM085601, AHB and TW006671, CA36622 CMI). We thank Jay Olsen for assistance with NMR, John N.A. Hooper for taxonomy discussions, and Lohi Matainaho for facilitating specimen collection in Papua New Guinea. The research reported in this publication utilized the High-Throughput Genomics Shared Resource at Huntsman Cancer Institute at the University of Utah and was supported by the National Cancer Institute of the National Institutes of Health under Award Number P30CA042014. The content is solely the responsibility of the authors and does not necessarily represent the official views of the NIH.

## Additional Files

Supplementary Figures

Supplementary Figures.pdf contains supplementary figures 1-5. Supplementary Figure 1 shows an image of the organism topsentinol L trisulfateTLT was isolated from. Supplementary Figure 2 shows the 1D NMR spectroscopy of halistanol sulfate and topsentinol L trisulfate in CD3OD. Supplementary Figure 3 shows the mass spectra for halistanol sulfate and topsentinol L trisulfate. Supplementary Figure 4 shows the 2D NMR of topenstinol L trisulfate in CD3OD. Supplementary Figure 5 shows that halistanol sulfate is not selective against BL-CL compared to luminal/HER2+ breast cancers. Supplementary Figure 6 demonstrates that the overexpression of AMPK and CHK1 sensitize BL-CL cell lines to topsentinol L trisulfate treatment.

Supplementary Information

Supplementary Information.doc contains additional details on methods used as well as the supplementary figure and table legends.

Supplementary Table 1

Supplementary Table 1.pdf contains a listing of all of the breast cancer cell lines used in this study along with ER, PR, HER2 status, gene-expression subtype, media in which it was maintained, and the ATCC number for the cell line.

Supplementary Table 2

Supplementary Table 2.pdf shows which cell lines were used in the different drug screens used to identify topsentinol L trisulfate.

Supplementary Table 3

Supplementary Table 3.pdf describes the chemical shifts in the NMR spectra used to distinguish topsentinol L trisulfate from halistanol sulfate.

Supplementary Table 4

Supplementary Table 4.pdf summarizes the reverse phase protein array (RPPA) results highlighting the proteins that were most significantly altered following TLT treatment for 6 hrs.

Supplementary Table 5

Supplementary Table 5.pdf shows that the EC50 to TLT decreases in breast cancer cell lines following transduction with AMPK and CHK1.

