## Supplementary Figures for "Topsentinol L Trisulfate, a new Marine Natural Product, that Targets Basal-like and Claudin-low Breast Cancers"

Supplementary Figure 1

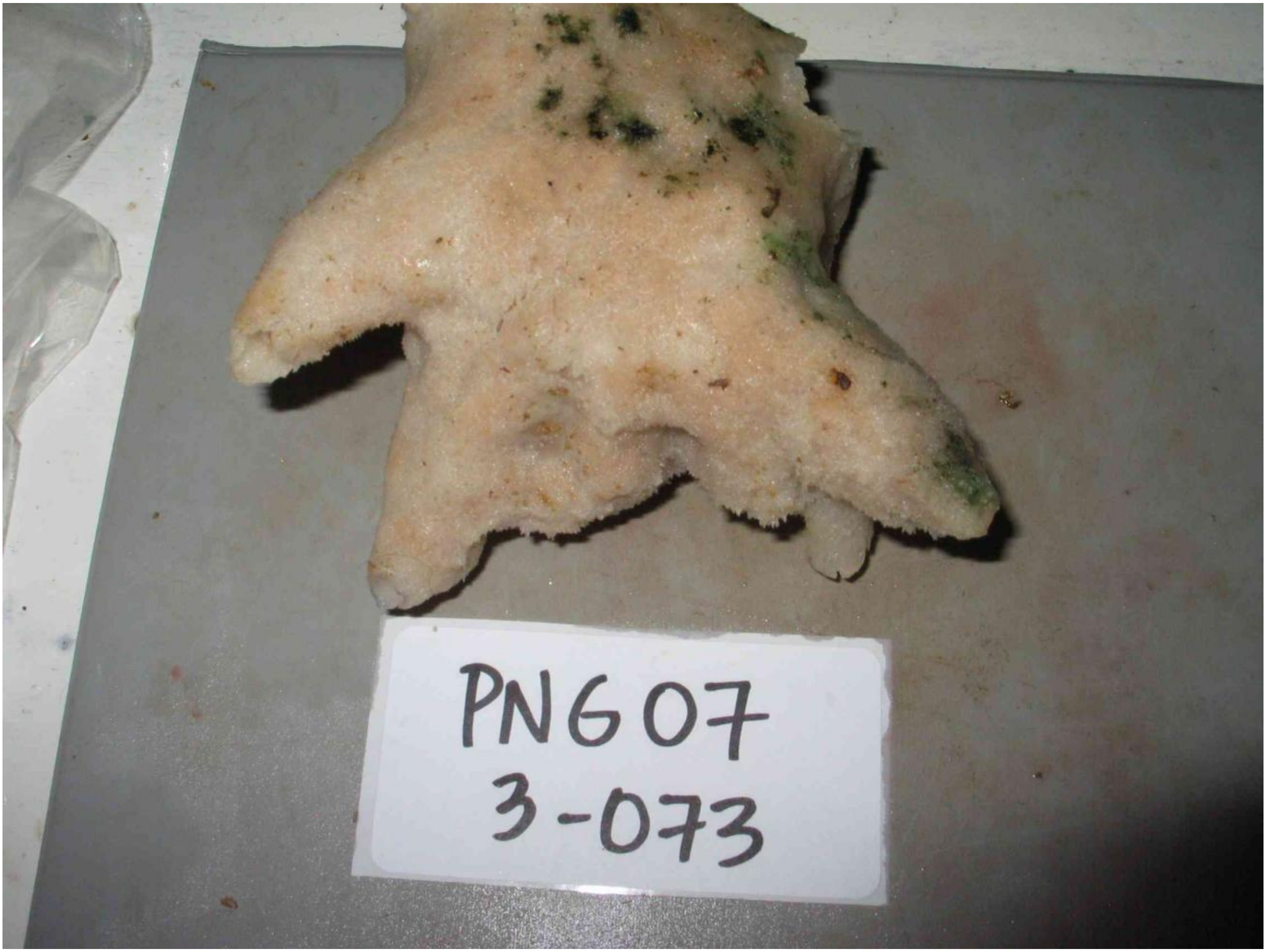

Supplementary Figure 2A

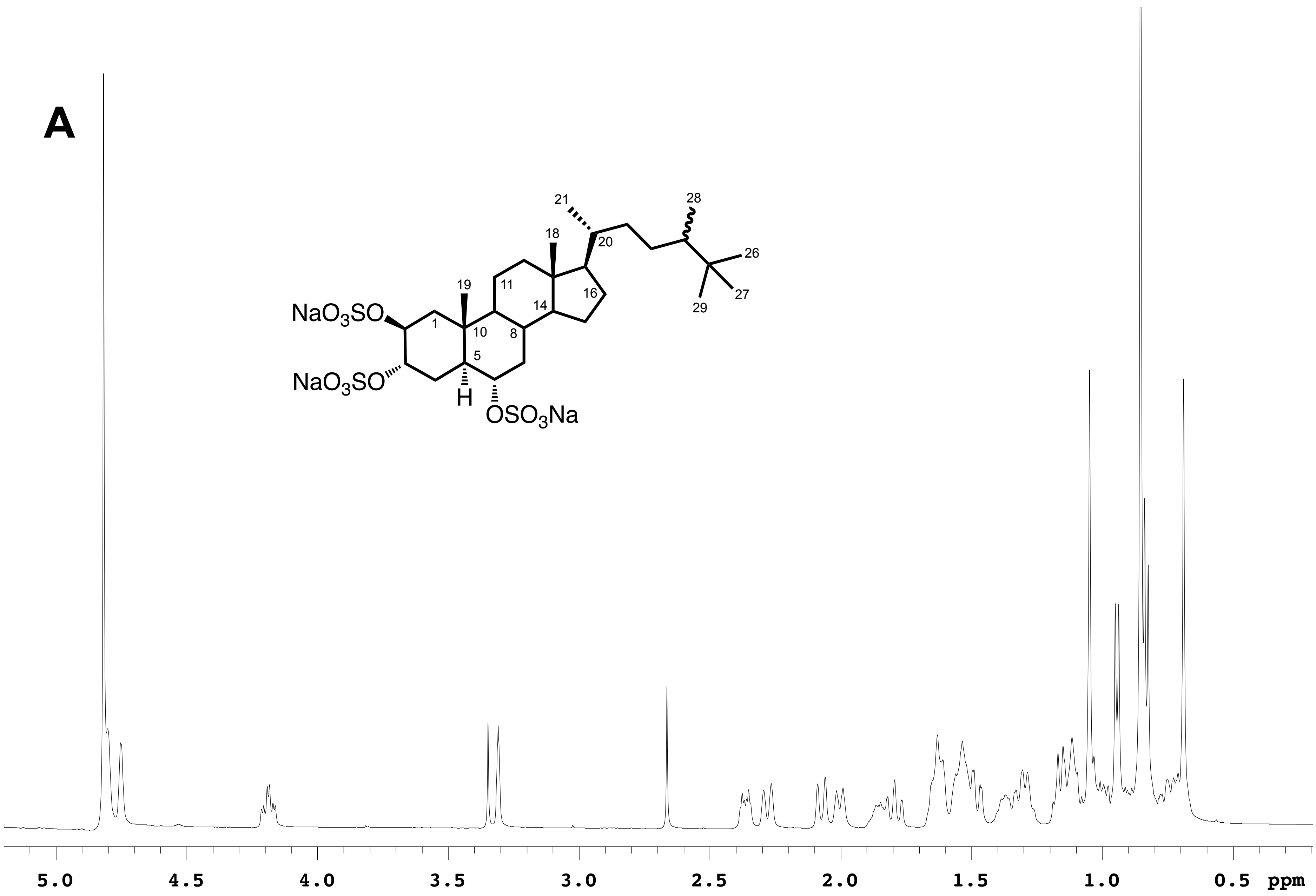

Supplementary Figure 2B

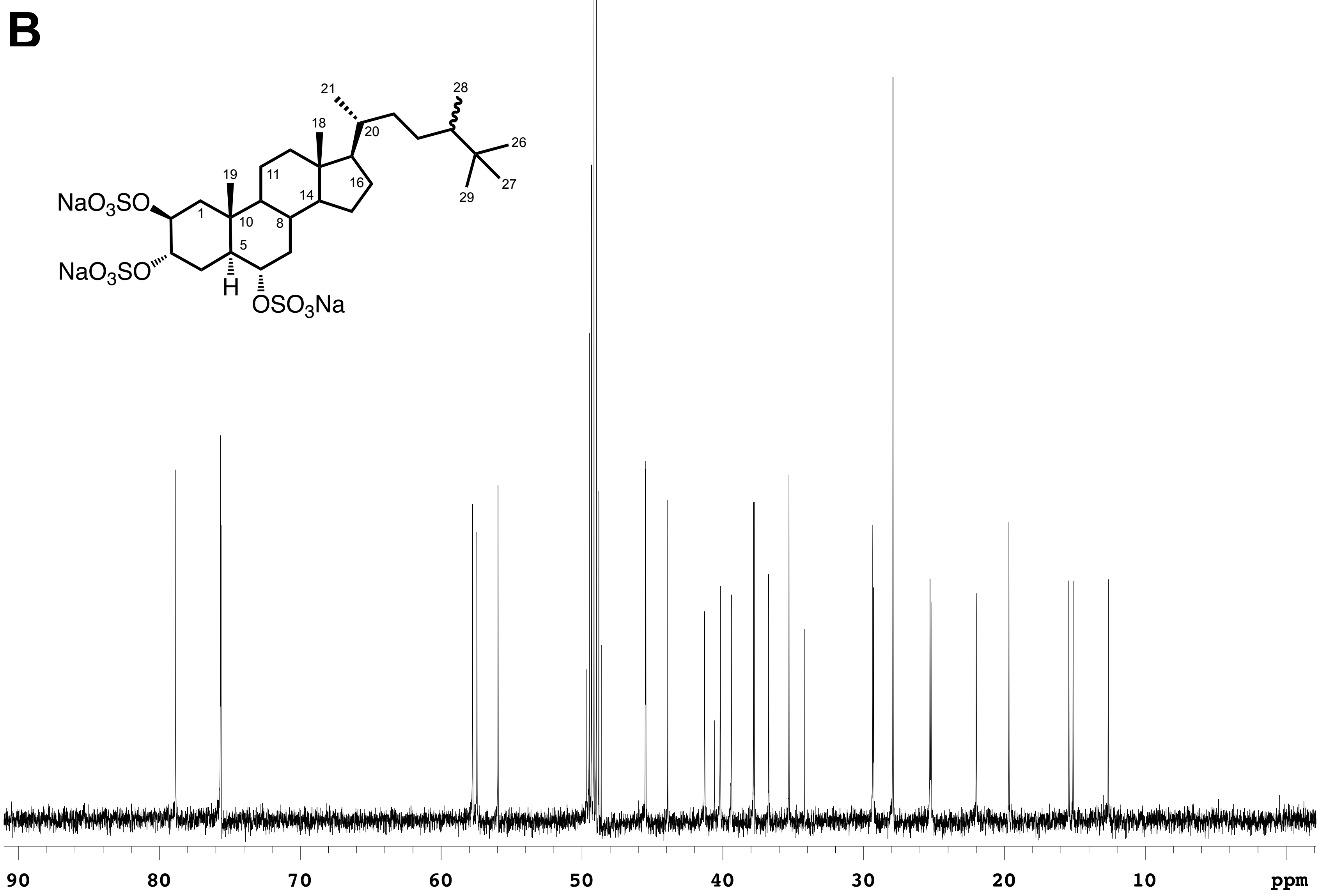

Supplementary Figure 2C

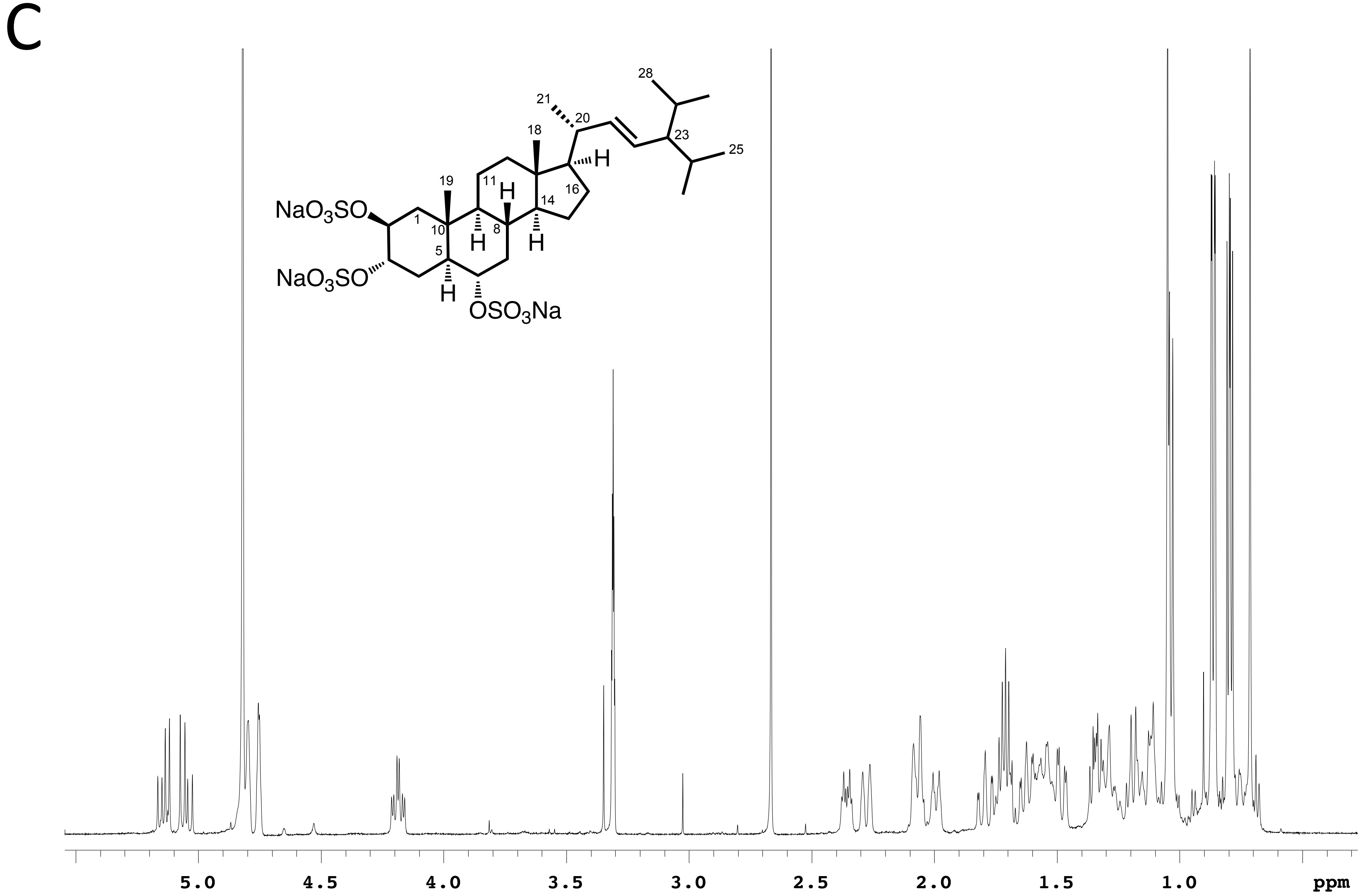

Supplementary Figure 2D

D

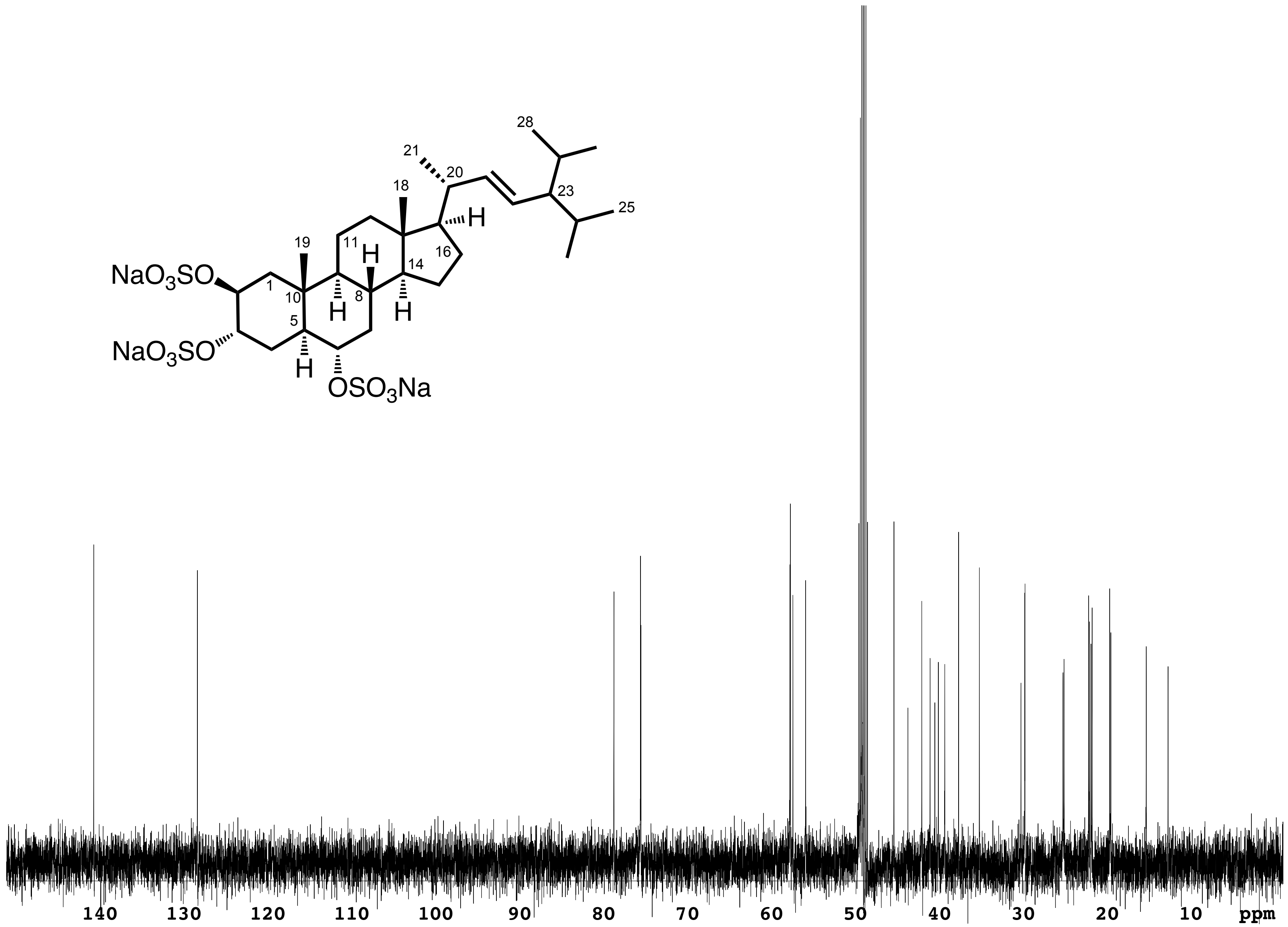

Supplementary Figure 3A

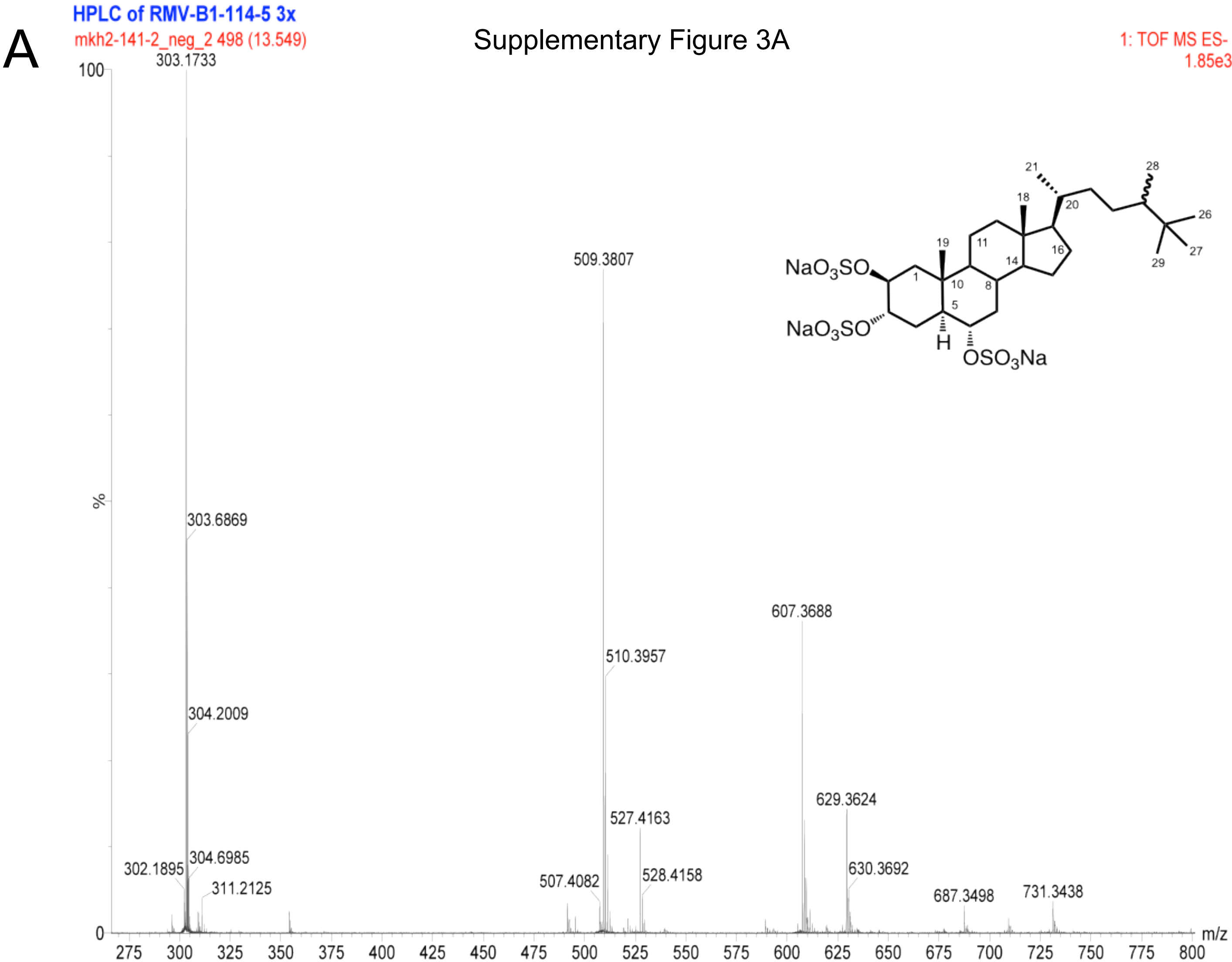

Supplementary Figure 3B

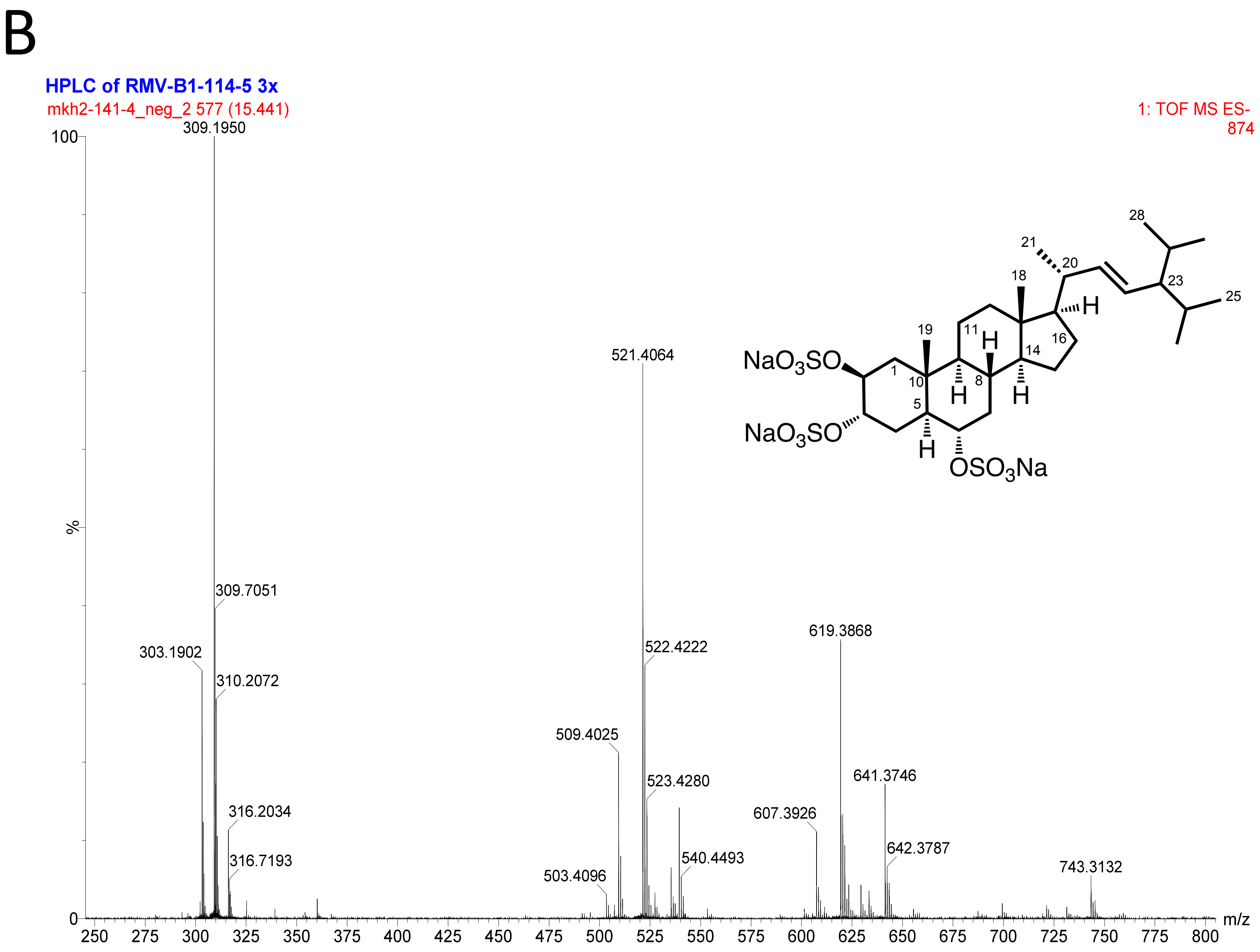

Supplementary Figure 4A

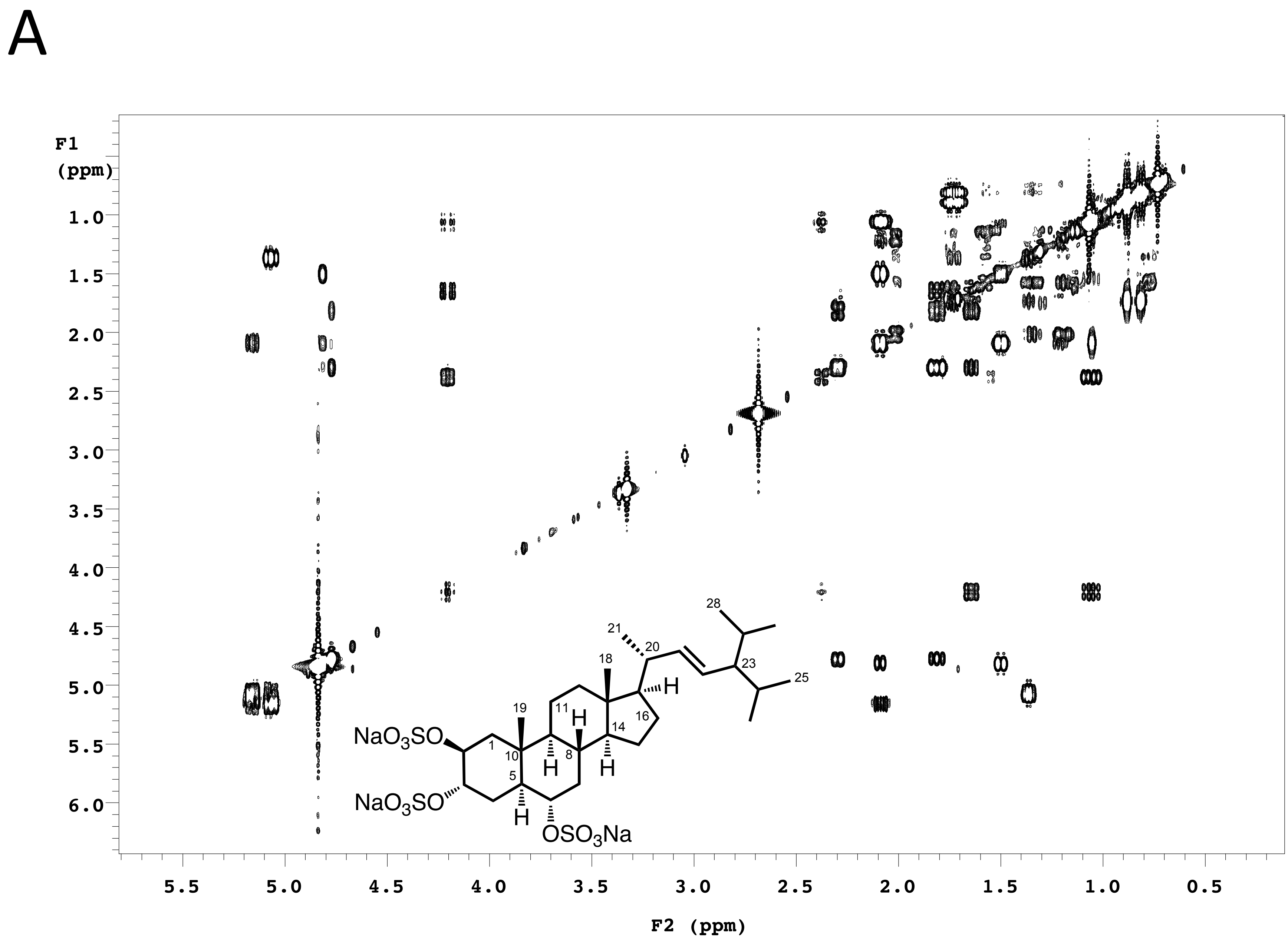

Supplementary Figure 4B

B

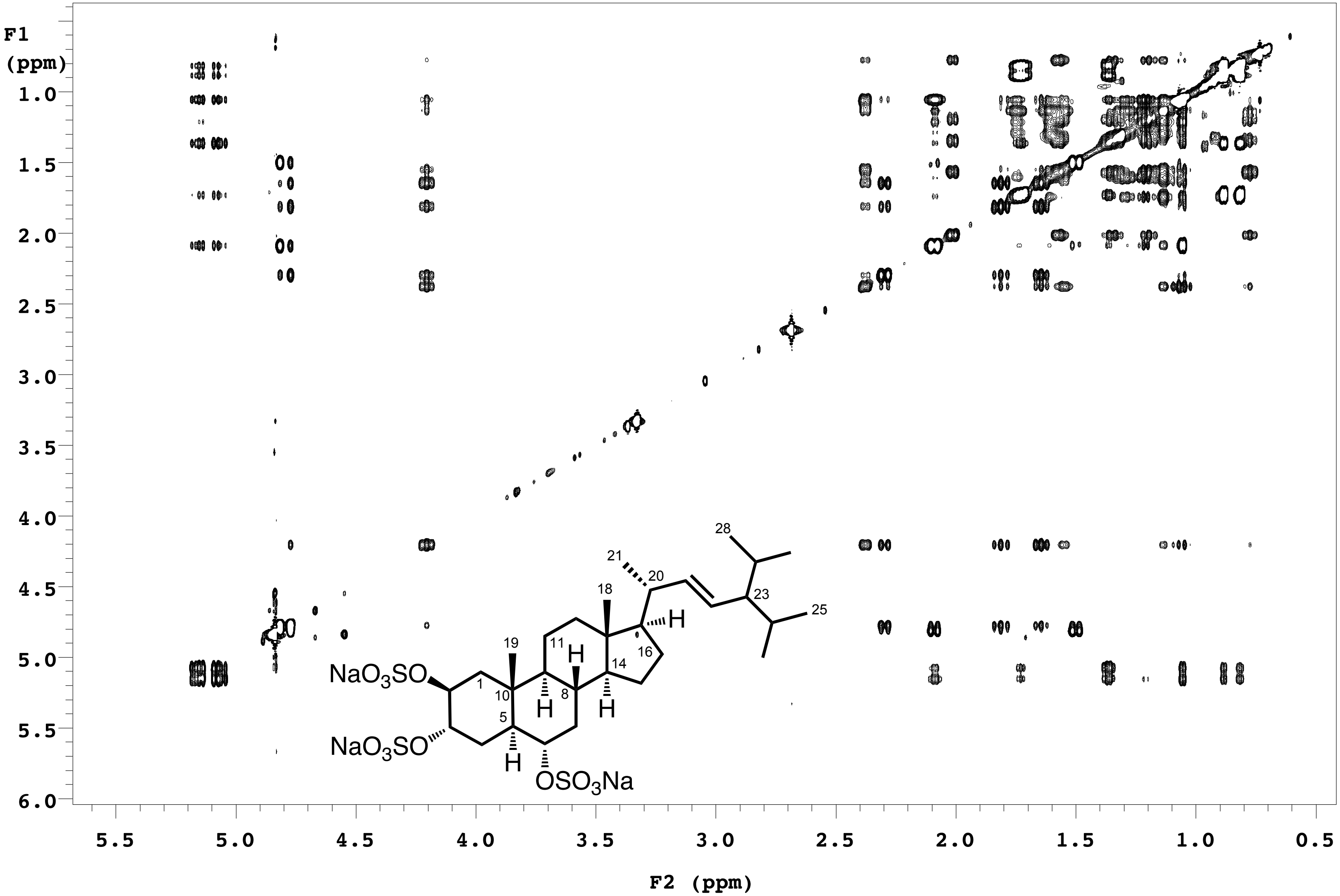

Supplementary Figure 4C

C

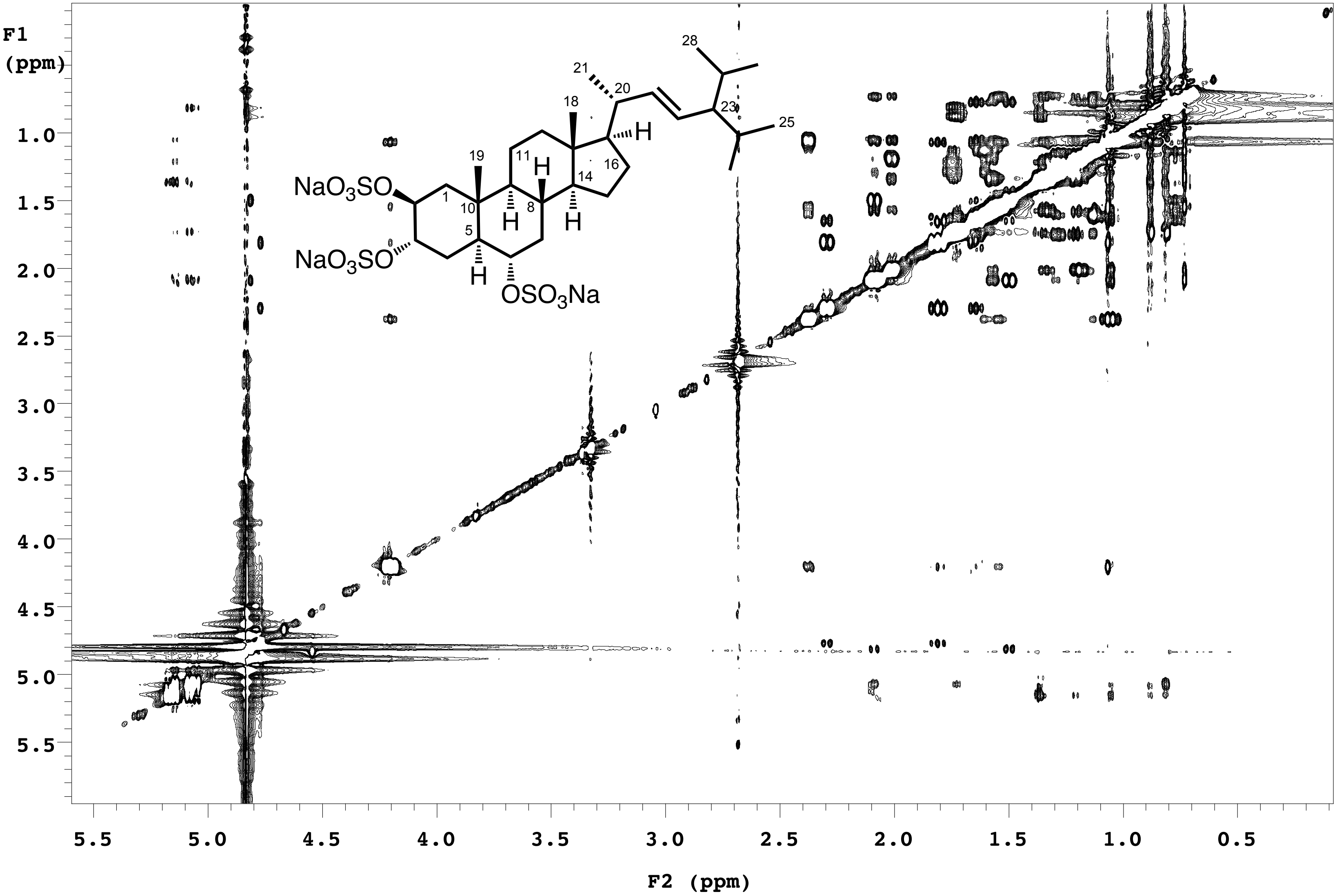

Supplementary Figure 4D

D

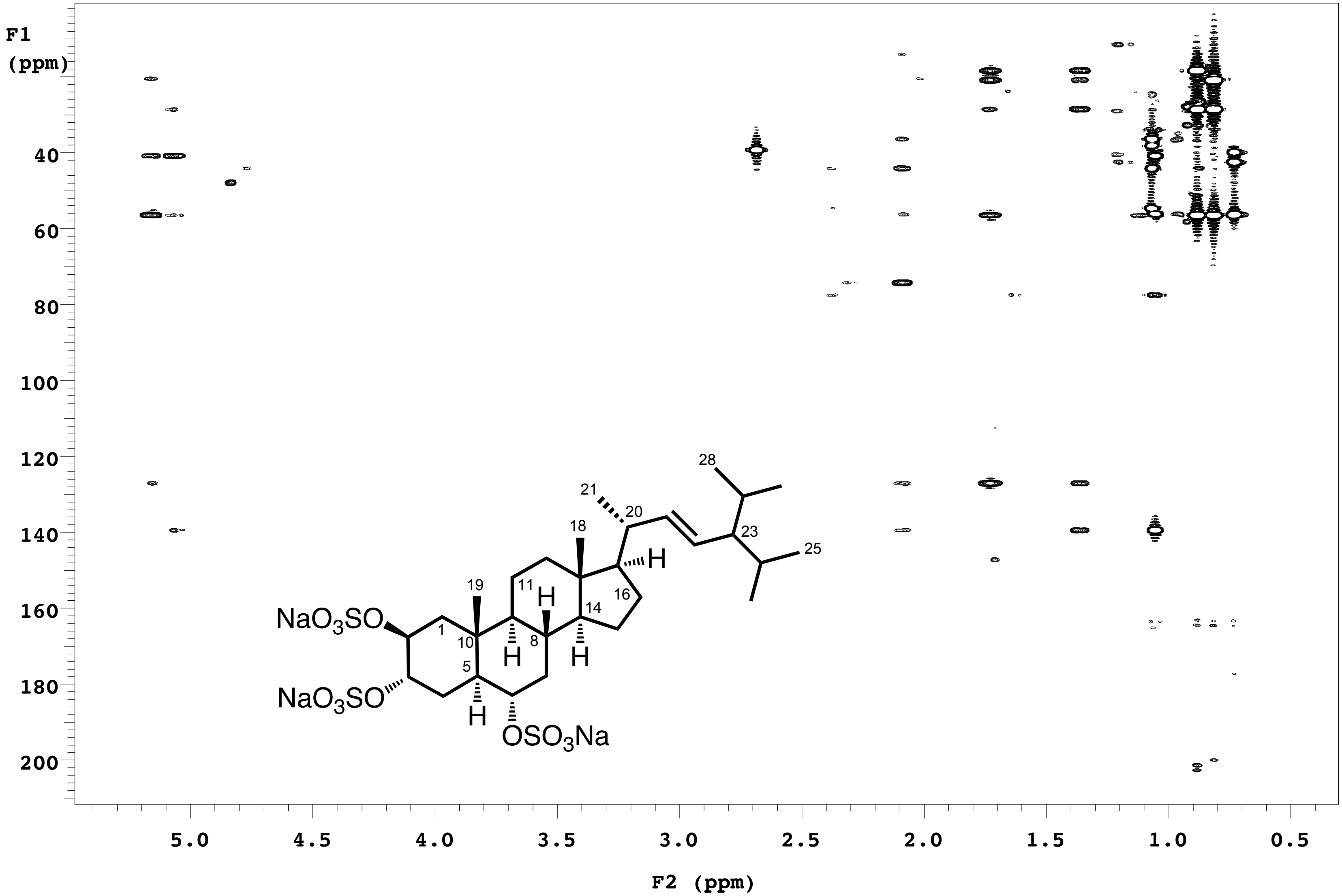

Supplementary Figure 4E

# E

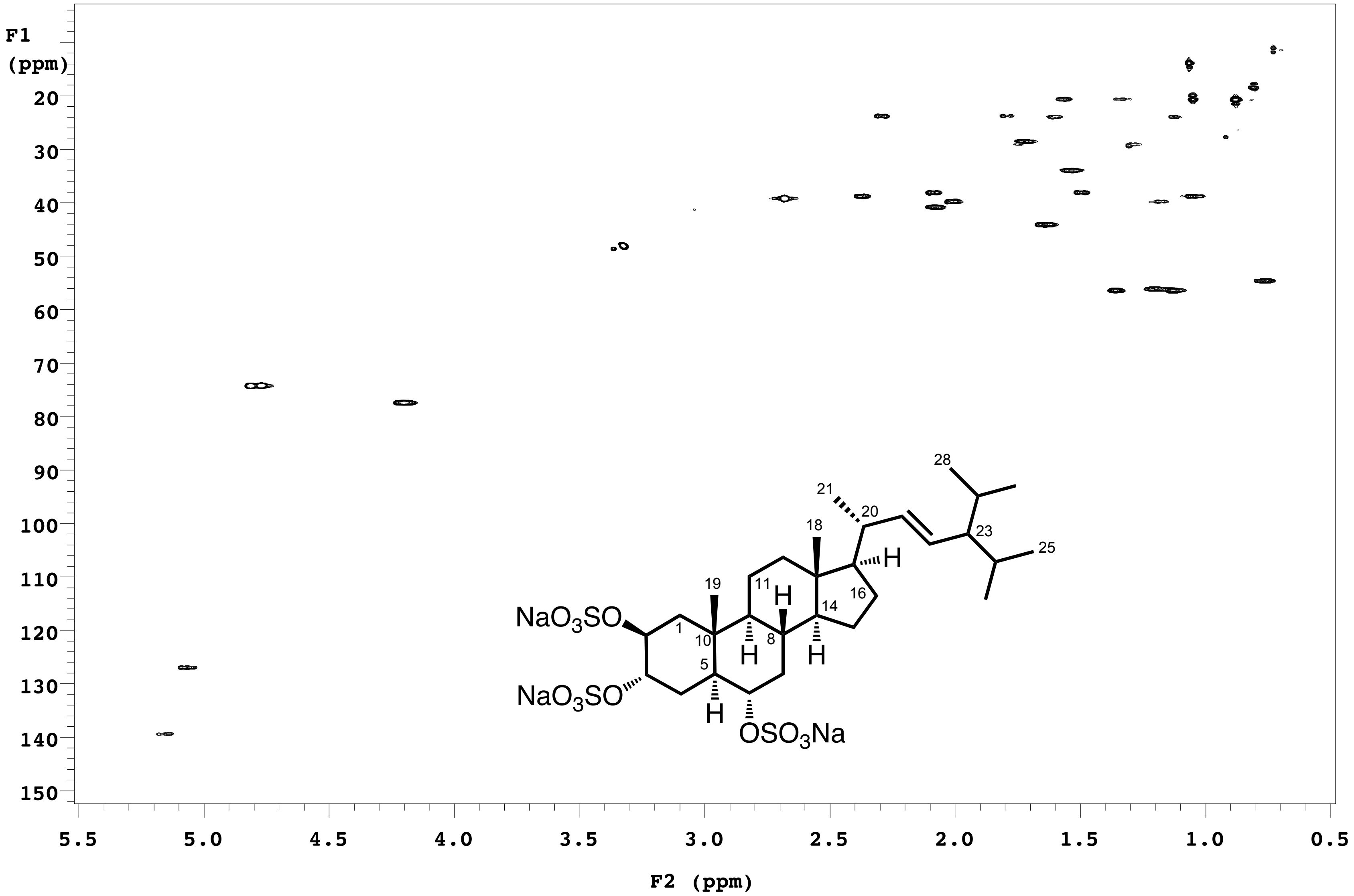

Supplementary Figure 5

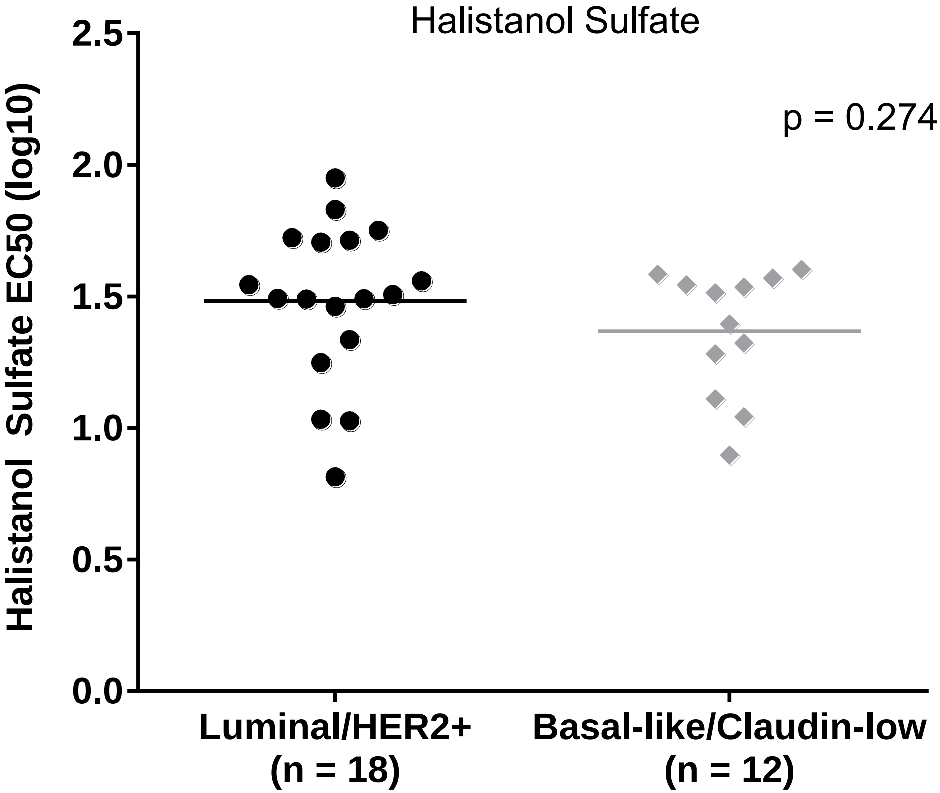

### Supplementary Figure 6

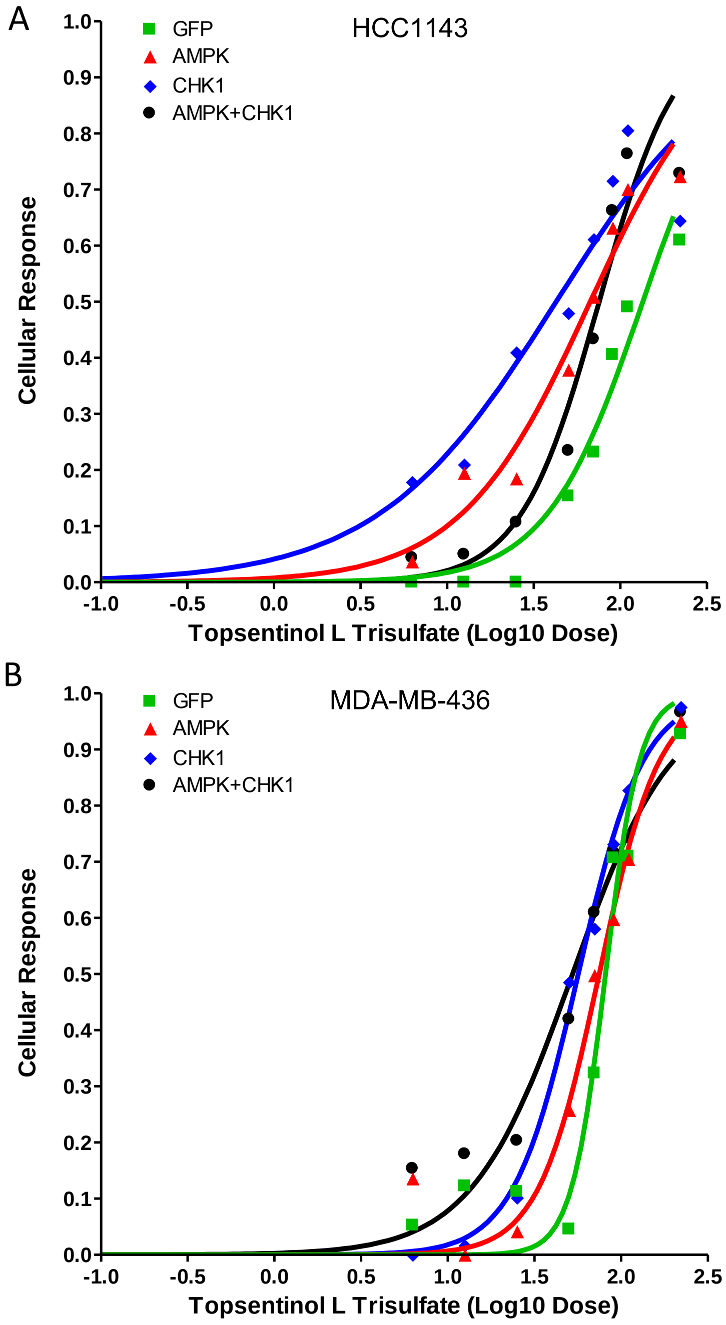
