## Supplementary Table 1 for "Topsentinol L Trisulfate, a new Marine Natural Product, that Targets Basal-like and Claudin-low Breast Cancers"

| Breast Cancer Cell Line | ER | PR | HER2 | Gene-expression Subtype | Media | ATCC Number |
|---|---|---|---|---|---|---|
| AU565 | - | - | + | luminal/HER2 positive | RPMI | CRL-2351 |
| BT-20 | - | - | - | basal | DMEM | HTB-19 |
| BT-474 | + | + | + | luminal/HER2 positive | DMEM | HTB-20 |
| BT-483 | + | + | - | luminal | RPMI | HTB-121 |
| BT-549 | - | - | - | claudin low | RPMI | HTB-122 |
| CAMA-1 | + | - | - | luminal | DMEM | HTB-21 |
| HCC1143 | - | - | - | basal | RPMI | CRL-2321 |
| HCC1395 | - | - | - | basal | RPMI | CRL-2324 |
| HCC1419 | - | - | + | luminal/HER2 positive | RPMI | CRL-2326 |
| HCC1500 | + | + | - | basal | RPMI | CRL-2329 |
| HCC1569 | - | - | + | basal/HER2 positive | RPMI | CRL-2330 |
| HCC1599 | - | - | - | basal | RPMI | CRL-2331 |
| HCC1806 | - | - | - | basal | RPMI | CRL-2335 |
| HCC1937 | - | - | - | basal | RPMI | CRL-2336 |
| HCC1954 | - | - | + | basal/HER2 positive | RPMI | CRL-2338 |
| HCC2218 | - | - | + | luminal | RPMI | CRL-2343 |
| HCC38 | - | - | - | claudin low | RPMI | CRL-2314 |
| HCC70 | - | - | - | basal | RPMI | CRL-2315 |
| Hs 578T | - | - | - | claudin low | DMEM | HTB-126 |
| JIMT-1 | - | - | + | basal/HER2 positive trastuzumab Res | DMEM | Unavailable |
| MCF7 | + | + | - | luminal | DMEM | HTB-22 |
| MDA-MB-134-VI | + | - | - | luminal | RPMI | HTB-23 |
| MDA-MB-157 | - | - | - | claudin low | DMEM | HTB-24 |
| MDA-MB-175-VII | + | - | - | luminal | RPMI | HTB-25 |
| MDA-MB-231 | - | - | - | claudin low | RPMI | HTB-26 |
| MDA-MB-415 | + | - | - | luminal | RPMI | HTB-128 |
| MDA-MB-436 | - | - | - | claudin low | RPMI | HTB-130 |
| MDA-MB-453 | - | - | - | luminal | RPMI | HTB-131 |
| MDA-MB-468 | - | - | - | basal | RPMI | HTB-132 |
| MDA-MB-361 | + | - | + | luminal/HER2 positive | DMEM | HTB-27 |
| SK-BR-3 | - | - | + | luminal/HER2 positive | RPMI | HTB-30 |
| T-47D | + | + | - | luminal | RPMI | HTB-133 |

| Breast Cancer Cell Line | ER | PR | HER2 | Gene-expression Subtype | Media | ATCC Number |
|---|---|---|---|---|---|---|
| UACC-812 | - | - | + | luminal/HER2 positive | DMEM | CRL-1897 |
| ZR-75-1 | + | - | - | luminal | RPMI | CRL-1500 |
| ZR-75-30 | + | - | + | luminal/HER2 positive | RPMI | CRL-1504 |

| Lung Cancer Cell Line | Subtype | K-Ras Mut | P53 Mut | EGFR Mut | Media | ATCC Number |
|---|---|---|---|---|---|---|
| A549 | Adenocarcinoma | G12S | x | x | F12K | CCL-185 |
| Calu-3 | Adenocarcinoma | x | M237I | x | EMEM | HTB-55 |
| NCI-H1155 | Large cell | Q61H | R273H | x | ACL-4 | CRL-5818 |
| NCI-H1355 | Adenocarcinoma | G13C | E285K | Q1159H | ACL-4 | CRL-5865 |
| NCI-H1373 | Adenocarcinoma | G12C | E339* | Intron | RPMI | CRL-5866 |
| NCI-H1395 | Adenocarcinoma | x | x | x | RPMI | CRL-5868 |
| NCI-H1437 | Adenocarcinoma | x | R267P | x | RPMI | CRL-5872 |
| NCI-H1563 | Adenocarcinoma | x | x | x | RPMI | CRL-5875 |
| NCI-H1581 | Large cell | x | Q144* | x | ACL-4 | CRL-5878 |
| NCI-H1650 | Adenocarcinoma | x | V225_splice | ELREA746del | RPMI | CRL-5883 |
| NCI-H1651 | Adenocarcinoma | x | C176Y | x | ACL-4 | CRL-5884 |
| NCI-H1693 | Adenocarcinoma | x | Q331_splice | x | RPMI | CRL-5887 |
| NCI-H1703 | Adenocarcinoma | x | A307_splice | x | RPMI | CRL-5889 |
| NCI-H1792 | Adenocarcinoma | G12C | E224_splice | x | RPMI | CRL-5895 |
| NCI-H1793 | Adenocarcinoma | x | R209* | C311F | HITES | CRL-5896 |
| NCI-H1944 | Adenocarcinoma | G13D | x | x | RPMI | CRL-5907 |
| NCI-H1975 | Adenocarcinoma | x | x | T790M, L858R | RPMI | CRL-5908 |
| NCI-H2009 | Adenocarcinoma | G12A | R273L | Intron | HITES | CRL-5911 |
| NCI-H2030 | Adenocarcinoma | G12C | G262V | x | RPMI | CRL-5914 |
| NCI-H2085 | Adenocarcinoma | x | ND | ND | ACL-4 | CRL-5921 |
| NCI-H2122 | Adenocarcinoma | G12C | C176F, Q16L | x | RPMI | CRL-5985 |
| NCI-H2126 | Adenocarcinoma | x | E62* | x | ACL-4 | CCL-256 |
| NCI-H23 | Adenocarcinoma | G12C | M246I | x | RPMI | CRL-5800 |
| NCI-H2405 | Adenocarcinoma | x | x | x | ACL-4 | CRL-5944 |
| NCI-H322 | Adenocarcinoma | x | R248L | x | RPMI | CRL-5806 |
| NCI-H358 | Adenocarcinoma | G12C | x | x | RPMI | CRL-5807 |
| NCI-H441 | Adenocarcinoma | G12V | R158L | x | RPMI | HTB-174 |

| Lung Cancer Cell Line | Subtype | K-Ras Mut | P53 Mut | EGFR Mut | Media | ATCC Number |
|---|---|---|---|---|---|---|
| NCI-H460 | Large cell | Q61H | x | x | RPMI | HTB-177 |
| NCI-H520 | Squamous | x | W146* | x | RPMI | HTB-182 |
| NCI-H522 | Adenocarcinoma | x | P191fs FRAME SHIFT DEL | x | RPMI | CRL-5810 |
| NCI-H661 | Large cell | x | R158L, S215I | x | RPMI | HTB-183 |
| NCI-H838 | Adenocarcinoma | x | x | x | RPMI | CRL-5844 |
| HCC4006 | Adenocarcinoma | x | x | ELR746del INFRAME | RPMI | CRL-2871 |
| SK-LU-1 | Adenocarcinoma | G12D | H193R | x | EMEM | HTB-57 |
| SK-MES-1 | Squamous | x | E298* | x | EMEM | HTB-58 |
| SW 1573 | Squamous | G12C | Intron | x | RPMI | CRL-2170 |
