## Supplementary Table 2 for "Topsentinol L Trisulfate, a new Marine Natural Product, that Targets Basal-like and Claudin-low Breast Cancers"

| **Breast Cancer Cell Line** | **Screen 1** | **Screen 2** | **Screen 3** | **Screen 4** |
|---|---|---|---|---|
| AU565 | | x | x | x |
| BT-20 | | x | x | x |
| BT-474 | x | x | x | x |
| BT-483 | | x | x | x |
| BT-549 | x | x | x | |
| CAMA-1 | | x | x | x |
| HCC1143 | | x | x | x |
| HCC1395 | | x | x | x |
| HCC1419 | | x | x | x |
| HCC1500 | | x | | |
| HCC1569 | | x | x | x |
| HCC1599 | | x | x | |
| HCC1806 | | x | x | x |
| HCC1937 | | x | x | x |
| HCC1954 | | x | x | x |
| HCC2218 | | x | x | x |
| HCC38 | | x | x | x |
| HCC70 | | x | x | x |
| Hs 578T | x | x | x | x |
| JIMT-1 | | x | x | x |
| MCF7 | x | x | x | x |
| MDA-MB-134-VI | | x | x | x |
| MDA-MB-157 | | x | x | x |
| MDA-MB-175-VII | | x | x | x |
| MDA-MB-231 | x | x | x | x |
| MDA-MB-415 | | x | x | x |
| MDA-MB-436 | | x | x | x |
| MDA-MB-453 | | x | x | x |
| MDA-MB-468 | | x | x | x |
| MDA-MB-361 | x | x | x | |
| SK-BR-3 | | x | x | x |
| T-47D | x | x | x | x |
| UACC-812 | | x | x | x |
| ZR-75-1 | | x | x | x |
| ZR-75-30 | | x | | |

| **Lung Cancer Cell Line** | **Screen 1** | **Screen 2** |
|---|---|---|
| A549 | | x |
| Calu-3 | | x |
| NCI-H1155 | | x |
| NCI-H1355 | | x |
| NCI-H1373 | | x |
| NCI-H1395 | | x |
| NCI-H1437 | | x |
| NCI-H1563 | x | x |
| NCI-H1581 | x | x |
| NCI-H1650 | x | x |
| NCI-H1651 | | x |
| NCI-H1693 | | x |
| NCI-H1703 | | x |
| NCI-H1792 | | x |
| NCI-H1793 | | x |
| NCI-H1944 | x | x |
| NCI-H1975 | x | x |
| NCI-H2009 | | x |
| NCI-H2030 | | x |
| NCI-H2085 | | x |
| NCI-H2122 | | x |
| NCI-H2126 | | x |
| NCI-H23 | x | x |
| NCI-H2405 | | x |
| NCI-H322 | | x |
| NCI-H358 | | x |
| NCI-H441 | | x |
| NCI-H460 | | x |
| NCI-H520 | x | x |
| NCI-H522 | | x |
| NCI-H661 | x | x |
| NCI-H838 | | x |
| HCC4006 | x | x |
| SK-LU-1 | | x |
| SK-MES-1 | | x |
| SW 1573 | | x |
