## Supplementary Table 3 for "Topsentinol L Trisulfate, a new Marine Natural Product, that Targets Basal-like and Claudin-low Breast Cancers"

| | Halistanol sulfate (mkh2-141-2) | | Topsentinol L trisulfate (mkh2-141-4) | |
|---|---|---|---|---|
| Position | $\delta_C$ (mult.) | $\delta_H$ (mult., $J$ in Hz) | $\delta_C$ (mult.) | $\delta_H$ (mult., $J$ in Hz) |
| 1 | 39.4 | 2.07 (br d, 14.3) | 39.4 | 2.07 (m) |
| | | 1.48 (dd, 15.4, 2.9) | | 1.48 (dd, 14.9, 3.7) |
| 2 | 75.7 | 4.80 (s) | 75.7 | 4.80 (s) |
| 3 | 75.6 | 4.75 (s) | 75.6 | 4.75 (br d, 22.3) |
| 4 | 25.2 | 2.28 (br d, 14.6) | 25.2 | 2.28 (br d, 14.5) |
| | | 1.79 (br t, 13.4) | | 1.78 (m) |
| 5 | 45.5 | 1.63 (m) | 45.5 | 1.62 (m) |
| 6 | 78.9 | 4.19 (td, 11.1, 4.4) | 78.9 | 4.19 (td, 11.1, 4.4) |
| 7 | 40.2 | 2.37 (dt, 12.2, 4.3) | 40.2 | 2.36 (dt, 12.1, 4.3) |
| | | 1.05 (m) | | 1.04 (m) |
| 8 | 35.3 | 1.53 (m) | 35.3 | 1.53 (m) |
| 9 | 56.0 | 0.76 (m) | 56.0 | 0.75 (m) |
| 10 | 37.8 | | 37.8 | |
| 11 | 22.0 | 1.54 (m) | 22.0 | 1.55 (m) |
| | | 1.31 (m) | | 1.31 (m) |
| 12 | 41.3 | 2.00 (br d, 12.4) | 41.2 | 2.00 (dt, 12.4, 2.9) |
| | | 1.15 (m) | | 1.17 (m) |
| 13 | 43.9 | | 43.8 | |
| 14 | 57.8 | 1.11 (m) | 57.8 | 1.11 (m) |
| 15 | 25.3 | 1.63 (m) | 25.3 | 1.59 (m) |
| | | 1.11 (m) | | 1.11 (m) |
| 16 | 29.3 | 1.86 (m) | 30.3 | 1.73 (m) |
| | | 1.29 (m) | | 1.29 (m) |
| 17 | 57.5 | 1.16 (m) | 57.5 | 1.19 (m) |
| 18 | 12.6 | 0.69 (s) | 12.8 | 0.71 (s) |
| 19 | 15.4 | 1.05 (s) | 15.4 | 1.05 (s) |
| 20 | 37.8 | 1.38 (m) | 42.2 | 2.06 (m) |
| 21 | 19.7 | 0.94 (d) | 21.9 | 1.05 (d, 7.2) |
| 22 | 36.8 | 1.56 (m) | 140.8 | 5.14 (dd, 15.2, 8.5) |
| | | 0.90 (m) | | NA |
| 23 | 29.4 | 1.63 (m) | 128.5 | 5.05 (dd, 15.6, 9.5) |
| | | 0.71 (m) | | NA |
| 24 | 45.5 | 0.99 (m) | 57.8 | 1.34 (m) |
| 25 | 34.2 | | 29.9 [a] | 1.70 (m) [a] |
| 26 | 27.9 | 0.86 (s) | 22.3 [a] | 0.86 (dd, 6.6, 1.6) [a] |
| 27 | 27.9 | 0.86 (s) | 19.8 [a] | 0.80 (dd, 6.7, 4.7) [a] |
| 28 | 15.1 | 0.83 (d) | 29.9 [b] | 1.70 (m) [b] |
| 29 | 27.9 | 0.86 (s) | 22.2 [b] | 0.86 (dd, 6.6, 1.6) [b] |
| 30 | NA | NA | 19.6 [b] | 0.80 (dd, 6.7, 4.7) [b] |

[a] Chemical shifts cannot be distinguished from those denoted by [b].
