## Supplementary Table 4 for "Topsentinol L Trisulfate, a new Marine Natural Product, that Targets Basal-like and Claudin-low Breast Cancers"

| Protein | Paired TTEST | DMSO Av | TLT Av | Mean diff DMSO-M4 | % Change |
|---|---|---|---|---|---|
| 14-3-3-beta | 0.297467434 | 0.1205898 | 0.17484 | -0.054254273 | 31.03008862 |
| 14-3-3-epsilon | 0.127055041 | 0.0924194 | 0.08764 | 0.004775493 | -5.448744719 |
| 14-3-3-zeta | 0.611359948 | 0.6997833 | 0.69274 | 0.007044538 | -1.016911279 |
| 4E-BP1 | 0.88102686 | 0.2872652 | 0.286 | 0.001265921 | -0.442630957 |
| 4E-BP1_pS65 | 0.688000793 | 0.6132918 | 0.60668 | 0.006608546 | -1.089290952 |
| 4E-BP1_pT37_T46 | 0.052849589 | 0.7249001 | 0.83875 | -0.11385355 | 13.57413446 |
| 53BP1 | 0.129295665 | 0.4471605 | 0.50738 | -0.060216947 | 11.86827429 |
| ACC_pS79 | 0.023981823 | 0.6414503 | 0.51904 | 0.122410764 | -23.5840916 |
| ACC1 | 0.057831968 | 0.7090081 | 0.66169 | 0.047320333 | -7.151459425 |
| ACVRL1 | 0.204285214 | 0.0932188 | 0.09812 | -0.004902111 | 4.995992268 |
| ADAR1 | 0.722298854 | 0.1220255 | 0.12328 | -0.001252066 | 1.015647984 |
| Akt | 0.019548209 | 0.572984 | 0.52345 | 0.04953045 | -9.462243974 |
| Akt_pS473 | 0.956418547 | 0.3849057 | 0.38686 | -0.001955655 | 0.505518217 |
| Akt_pT308 | 0.848831423 | 1.0898163 | 1.07076 | 0.019051852 | -1.779275781 |
| AMPK-alpha | 0.494529968 | 0.8115008 | 0.82971 | -0.01821201 | 2.194977551 |
| AMPK-alpha_pT172 | 0.030926158 | 0.2468539 | 0.18237 | 0.06448569 | -35.360163 |
| Annexin-I | 0.202592026 | 0.2944506 | 0.35136 | -0.056913782 | 16.19793713 |
| Annexin-VII | 0.479745507 | 0.1808896 | 0.19652 | -0.015631946 | 7.954316733 |
| AR | 0.086105524 | 0.1613788 | 0.15309 | 0.008291424 | -5.416139865 |
| A-Raf | 0.021802658 | 0.3242939 | 0.30103 | 0.023259764 | -7.726620108 |
| ARHI | 0.374227021 | 0.0561487 | 0.0545 | 0.001644846 | -3.017852599 |
| ATM | 0.310104666 | 0.1761461 | 0.17153 | 0.004614255 | -2.690027967 |
| ATM_pS1981 | 0.937022455 | 0.0777696 | 0.07816 | -0.000390239 | 0.499282649 |
| ATP5H | 0.186610373 | 0.0743909 | 0.07776 | -0.003372651 | 4.337059112 |
| ATR | 0.92837797 | 0.2607643 | 0.26007 | 0.000696483 | -0.26780838 |
| Bad_pS112 | 0.010420165 | 0.4945067 | 0.55515 | -0.060640319 | 10.9232895 |
| Bak | 0.008721632 | 0.1909752 | 0.2054 | -0.0144239 | 7.022377178 |
| BAP1 | 0.259977565 | 0.2223549 | 0.21511 | 0.007244081 | -3.367604315 |
| Bax | 0.343854359 | 0.3507281 | 0.36731 | -0.016583733 | 4.514891461 |
| b-Catenin | 0.584632507 | 0.6792402 | 0.71601 | -0.036766439 | 5.134930087 |
| b-Catenin_pT41_S45 | 0.070481607 | 0.1303322 | 0.13996 | -0.009625835 | 6.877656406 |
| Bcl2 | 0.422689711 | 0.0737728 | 0.07546 | -0.001686947 | 2.235559768 |
| Bcl-xL | 0.13933645 | 0.5464368 | 0.57938 | -0.032944392 | 5.68613386 |
| Beclin | 0.302512426 | 0.1887283 | 0.22117 | -0.032440994 | 14.66794427 |
| Bid | 0.654051893 | 0.2214305 | 0.22455 | -0.003117031 | 1.388138801 |
| Bim | 0.601368016 | 0.5163407 | 0.53035 | -0.014011711 | 2.641962346 |
| B-Raf | 0.032395628 | 1.009608 | 0.94963 | 0.059981572 | -6.316333569 |
| B-Raf_pS445 | 0.778696211 | 0.1716865 | 0.17297 | -0.001285076 | 0.742940326 |
| BRCA2 | 0.634796435 | 0.1778776 | 0.18588 | -0.008002506 | 4.305198123 |
| Caspase-7-cleaved | 0.199719195 | 0.0678278 | 0.07255 | -0.004718936 | 6.504683391 |
| Caspase-8 | 0.111844814 | 0.3088891 | 0.29177 | 0.017117839 | -5.866869961 |
| Caveolin-1 | 0.330102248 | 0.454963 | 0.44035 | 0.014615083 | -3.31898519 |
| CD29 | 0.347436034 | 0.031597 | 0.03009 | 0.001504719 | -5.00034688 |
| CD31 | 0.027495297 | 0.0240145 | 0.02141 | 0.002608377 | -12.18521243 |

| Protein | Paired TTEST | DMSO Av | TLT Av | Mean diff DMSO-M4 | % Change |
|---|---|---|---|---|---|
| CD49b | 0.000681184 | 0.090206 | 0.08321 | 0.006999799 | -8.412591597 |
| CDK1 | 0.527951409 | 0.2966172 | 0.30303 | -0.006412135 | 2.116011117 |
| Chk1 | 0.517023385 | 0.4436631 | 0.43806 | 0.005598172 | -1.277931995 |
| Chk1_pS345 | 0.012431443 | 0.2876721 | 0.24789 | 0.039780631 | -16.0476008 |
| Chk2 | 0.900777361 | 0.3054057 | 0.30321 | 0.002196345 | -0.724366013 |
| Chk2_pT68 | 0.234845145 | 0.1516815 | 0.15712 | -0.005434188 | 3.458717341 |
| c-Jun_pS73 | 0.039255625 | 0.2882244 | 0.34707 | -0.058847805 | 16.95549391 |
| c-Kit | 0.137841447 | 0.1486842 | 0.15822 | -0.009531557 | 6.024403154 |
| Claudin-7 | 0.838258667 | 0.5496062 | 0.53524 | 0.014368512 | -2.684510622 |
| c-Met | 0.100343269 | 0.149599 | 0.14234 | 0.007259813 | -5.100362748 |
| c-Met_pY1234_Y1235 | 0.607157419 | 0.1791378 | 0.1818 | -0.002660777 | 1.463584969 |
| c-Myc | 0.823539509 | 0.3123089 | 0.3083 | 0.00400808 | -1.300054814 |
| Collagen-VI | 0.581827632 | 0.1085949 | 0.09859 | 0.010000297 | -10.14284249 |
| Complex-II-Subunit | 0.019357592 | 0.4883395 | 0.5321 | -0.043759206 | 8.223889209 |
| Cox2 | 0.267016058 | 0.1049197 | 0.17838 | -0.073460635 | 41.18203035 |
| Cox-IV | 0.21180139 | 0.0470478 | 0.04401 | 0.003040435 | -6.908917587 |
| C-Raf | 0.00569841 | 0.2922268 | 0.26106 | 0.031170474 | -11.94013341 |
| C-Raf_pS338 | 0.181233222 | 0.4905809 | 0.45703 | 0.033551782 | -7.341279215 |
| Cyclin-B1 | 0.047649153 | 1.5024802 | 1.70111 | -0.19862495 | 11.67622972 |
| Cyclin-D1 | 0.255149632 | 0.3947857 | 0.37585 | 0.018939256 | -5.039094233 |
| Cyclin-E1 | 0.471229306 | 0.3046325 | 0.2948 | 0.009828166 | -3.333793248 |
| Cyclophilin-F | 0.588089846 | 1.2412809 | 1.12366 | 0.117620772 | -10.4676463 |
| DJ1 | 0.057596359 | 0.3065183 | 0.31909 | -0.012574353 | 3.940658557 |
| Dvl3 | 0.305796393 | 0.3631395 | 0.35437 | 0.008770662 | -2.475009209 |
| E2F1 | 0.465992121 | 0.0490497 | 0.05045 | -0.001399862 | 2.774776277 |
| E-Cadherin | 0.414543231 | 0.2309039 | 0.26116 | -0.030252555 | 11.58407426 |
| eEF2 | 0.024329272 | 0.4482607 | 0.42437 | 0.023888806 | -5.629214761 |
| eEF2K | 0.019692069 | 0.5732332 | 0.5177 | 0.055530627 | -10.72635653 |
| EGFR | 0.694306782 | 0.6800515 | 0.66891 | 0.011142733 | -1.665807521 |
| EGFR_pY1068 | 0.28897535 | 0.0780788 | 0.09203 | -0.013946575 | 15.15514849 |
| EGFR_pY1173 | 0.30499704 | 0.1638452 | 0.17395 | -0.010102281 | 5.80766297 |
| eIF4E | 0.125804471 | 0.8195001 | 0.79346 | 0.026037641 | -3.28152156 |
| eIF4G | 0.878724419 | 1.035565 | 1.03183 | 0.003731644 | -0.361651783 |
| ER-alpha | 0.504178772 | 0.0388962 | 0.0377 | 0.001192711 | -3.163397879 |
| ER-alpha_pS118 | 0.634911633 | 0.3953871 | 0.39946 | -0.004070958 | 1.019120408 |
| ERCC1 | 0.007891293 | 0.1646311 | 0.19196 | -0.02732517 | 14.23510243 |
| Ets-1 | 0.436109849 | 0.2073012 | 0.20972 | -0.002419888 | 1.153860178 |
| FAK | 0.00596371 | 0.3746999 | 0.35153 | 0.023166155 | -6.590023246 |
| FAK_pY397 | 0.080659613 | 0.0997996 | 0.08071 | 0.019094485 | -23.65957935 |
| FASN | 0.010279931 | 0.7383777 | 0.68099 | 0.057383377 | -8.426410979 |
| Fibronectin | 0.287511878 | 0.0348751 | 0.03279 | 0.002080442 | -6.34384711 |
| FoxM1 | 0.021099513 | 0.466201 | 0.42671 | 0.039490719 | -9.254691723 |
| FoxO3a | 0.551797236 | 0.1353121 | 0.1374 | -0.002085824 | 1.518089719 |
| FoxO3a_pS318_S321 | 0.943894769 | 0.5589604 | 0.56027 | -0.001312249 | 0.234216082 |
| G6PD | 0.228099421 | 0.0796185 | 0.07617 | 0.003448534 | -4.52742041 |

| Protein | Paired TTEST | DMSO Av | TLT Av | Mean diff DMSO-M4 | % Change |
|---|---|---|---|---|---|
| Gab2 | 0.129420955 | 0.2553089 | 0.24282 | 0.012488801 | -5.143232028 |
| GAPDH | 0.076587901 | 0.511228 | 0.41902 | 0.092207104 | -22.00536928 |
| GATA3 | 0.444543139 | 0.2890905 | 0.27977 | 0.009317527 | -3.330389022 |
| GCN5L2 | 0.581933997 | 0.4024414 | 0.41069 | -0.008251661 | 2.009204067 |
| GPBB | 0.693848285 | 0.2242394 | 0.23134 | -0.00710235 | 3.070067796 |
| GSK-3ab | 0.107780308 | 0.8094326 | 0.82645 | -0.017016942 | 2.059041866 |
| GSK-3ab_pS21_S9 | 0.550735827 | 0.6241206 | 0.64794 | -0.023816819 | 3.675790205 |
| GSK-3b_pS9 | 0.177310106 | 0.4761323 | 0.5083 | -0.03216499 | 6.327988207 |
| Gys | 0.995834943 | 0.518761 | 0.51881 | -5.10903E-05 | 0.009847553 |
| Gys_pS641 | 0.541992189 | 0.3732588 | 0.36622 | 0.007043452 | -1.923308627 |
| HER2 | 0.263192338 | 0.0281805 | 0.02605 | 0.002128034 | -8.16827614 |
| HER2_pY1248 | 0.104377998 | 0.0403153 | 0.04791 | -0.0075911 | 15.8456832 |
| HER3 | 0.192076665 | 0.2256087 | 0.20685 | 0.018756371 | -9.06751822 |
| HER3_pY1289 | 0.334961404 | 0.3471657 | 0.33777 | 0.009395071 | -2.781494189 |
| Heregulin | 0.356306262 | 0.158463 | 0.16414 | -0.005672826 | 3.456177062 |
| HIAP | 0.418473854 | 0.4569881 | 0.44928 | 0.00770568 | -1.715108173 |
| Histone-H3 | 0.859920263 | 0.1358833 | 0.13816 | -0.002276057 | 1.647415041 |
| IGF1R-beta | 0.633604069 | 0.2032754 | 0.20619 | -0.002917845 | 1.415102345 |
| IGFBP2 | 0.004643877 | 0.1284908 | 0.19396 | -0.065470005 | 33.7542394 |
| INPP4b | 0.224786181 | 0.1661943 | 0.16219 | 0.004006157 | -2.470067174 |
| IRS1 | 0.149211343 | 0.6053276 | 0.58946 | 0.015868518 | -2.692047293 |
| JAB1 | 0.551441944 | 0.0690752 | 0.06787 | 0.001201038 | -1.769508052 |
| JNK_pT183_Y185 | 0.622517253 | 0.1801214 | 0.18676 | -0.006641022 | 3.555867298 |
| JNK2 | 0.620583516 | 0.4413883 | 0.43619 | 0.005199641 | -1.192062542 |
| Lck | 0.404041823 | 0.1615087 | 0.15269 | 0.008820444 | -5.776766198 |
| MAPK_pT202_Y204 | 0.063897497 | 0.3657461 | 0.27199 | 0.09375541 | -34.47007848 |
| Mcl-1 | 0.869330062 | 0.3021391 | 0.30475 | -0.002608929 | 0.856093758 |
| MDM2_pS166 | 0.184798884 | 0.4228332 | 0.43978 | -0.016949566 | 3.854077221 |
| MEK1 | 0.749093243 | 0.4931058 | 0.48759 | 0.005517333 | -1.131555193 |
| MEK1_pS217_S221 | 0.330871457 | 0.4264982 | 0.40648 | 0.020019075 | -4.924994372 |
| MEK2 | 0.046725637 | 0.262228 | 0.24935 | 0.012881695 | -5.166185233 |
| Merlin | 0.110425164 | 0.6224777 | 0.57406 | 0.048420054 | -8.434701674 |
| MIG6 | 0.65818707 | 0.2714421 | 0.26646 | 0.004981295 | -1.869428612 |
| MSH2 | 0.232864132 | 0.7096262 | 0.69032 | 0.019305584 | -2.796611216 |
| MSH6 | 0.353124034 | 0.8008488 | 0.8208 | -0.019953461 | 2.43097042 |
| mTOR | 0.10247673 | 1.0000496 | 0.97459 | 0.025457724 | -2.612141883 |
| mTOR_pS2448 | 0.535634771 | 0.5736505 | 0.58052 | -0.006872585 | 1.183860668 |
| Myosin-11 | 0.130662479 | 0.2516351 | 0.24104 | 0.010593952 | -4.3950794 |
| Myosin-IIa_pS1943 | 0.136330833 | 0.5590272 | 0.52052 | 0.038505586 | -7.397499919 |
| NAPSIN-A | 0.000244622 | 0.1888714 | 0.1789 | 0.009967334 | -5.571328945 |
| N-Cadherin | 0.115979527 | 0.108239 | 0.0995 | 0.008736554 | -8.780236006 |
| NDRG1_pT346 | 0.168014205 | 0.3168807 | 0.37585 | -0.05897111 | 15.68998963 |
| NF-kB-p65_pS536 | 0.700093069 | 0.5674735 | 0.54107 | 0.026404604 | -4.880082075 |
| Notch1 | 0.661226615 | 0.6396916 | 0.65434 | -0.01464397 | 2.237990814 |
| N-Ras | 0.424040717 | 0.0373128 | 0.03559 | 0.001720188 | -4.83299624 |

| Protein | Paired TTEST | DMSO Av | TLT Av | Mean diff DMSO-M4 | % Change |
|---|---|---|---|---|---|
| p16INK4a | 0.089057378 | 0.2589195 | 0.24097 | 0.017949144 | -7.448692453 |
| p21 | 0.072152138 | 0.589791 | 0.68726 | -0.09746523 | 14.18179013 |
| p27_pT157 | 0.628732156 | 0.4132618 | 0.42135 | -0.008085064 | 1.918861914 |
| p27_pT198 | 0.015523921 | 0.380493 | 0.4207 | -0.040209916 | 9.557793229 |
| p27-Kip-1 | 0.101448071 | 0.1353413 | 0.12438 | 0.010959469 | -8.811151583 |
| p38 | 0.03267437 | 0.8570567 | 0.8278 | 0.029257917 | -3.534423672 |
| p38_pT180_Y182 | 0.002755169 | 0.4029123 | 0.52386 | -0.120951629 | 23.08836548 |
| p38-alpha | 0.00774209 | 0.3541716 | 0.32024 | 0.033932183 | -10.59587949 |
| p53 | 0.314234566 | 0.0975719 | 0.09485 | 0.002724253 | -2.872240696 |
| p70-S6K_pT389 | 0.031044202 | 0.1669332 | 0.14084 | 0.026097196 | -18.53020434 |
| p70-S6K1 | 0.172006967 | 0.2027094 | 0.1965 | 0.006210831 | -3.160751165 |
| PAI-1 | 0.360871698 | 0.7728393 | 0.79675 | -0.02391276 | 3.001280111 |
| PARP1 | 0.967407249 | 2.6880892 | 2.69449 | -0.006399885 | 0.237517567 |
| PARP-cleaved | 0.118830095 | 0.0705842 | 0.06733 | 0.00325635 | -4.836555963 |
| Paxillin | 0.906536745 | 0.5151816 | 0.51694 | -0.001762799 | 0.341003507 |
| PCNA | 0.06253059 | 0.2458135 | 0.23039 | 0.015419488 | -6.692659587 |
| Pdcd-1L1 | 0.112960284 | 0.1537748 | 0.21549 | -0.061714605 | 28.63927851 |
| Pdcd4 | 0.2306148 | 0.4398906 | 0.47403 | -0.034134929 | 7.201073805 |
| PDGFR-beta | 0.638238272 | 0.7599954 | 0.79413 | -0.034136002 | 4.298533004 |
| PDK1 | 0.004328426 | 0.1989206 | 0.17397 | 0.024949699 | -14.34130609 |
| PDK1_pS241 | 0.010023724 | 0.4803055 | 0.44315 | 0.037151425 | -8.38341026 |
| PEA-15 | 0.575925151 | 0.4603699 | 0.4734 | -0.013030284 | 2.752488295 |
| PEA-15_pS116 | 0.07112505 | 0.4775941 | 0.51308 | -0.035485327 | 6.91614747 |
| PI3K-p110-alpha | 0.045315699 | 0.3424673 | 0.32277 | 0.019699858 | -6.103421879 |
| PI3K-p85 | 0.209222646 | 0.2749676 | 0.27946 | -0.004493387 | 1.607876193 |
| PKC-alpha | 0.896269303 | 0.7304674 | 0.7277 | 0.002768541 | -0.380451526 |
| PKC-alpha_pS657 | 0.795719345 | 0.4284552 | 0.43106 | -0.002602363 | 0.603716011 |
| PKC-beta-II_pS660 | 0.425784911 | 0.7686049 | 0.80879 | -0.040182907 | 4.968287878 |
| PKC-delta_pS664 | 0.045976568 | 0.3448394 | 0.36918 | -0.024340036 | 6.593008843 |
| PMS2 | 0.025956644 | 0.801951 | 0.74703 | 0.054920915 | -7.351901528 |
| Porin | 0.243883965 | 0.0857229 | 0.08904 | -0.003322 | 3.730702144 |
| PR | 0.663238212 | 0.1169763 | 0.11619 | 0.000789801 | -0.679769967 |
| PRAS40_pT246 | 0.365275156 | 0.8290985 | 0.86475 | -0.035650351 | 4.122624665 |
| PREX1 | 0.101304994 | 0.1049792 | 0.09812 | 0.006861336 | -6.992955773 |
| PTEN | 0.782392774 | 0.1851321 | 0.18314 | 0.001988066 | -1.085520762 |
| Rab11 | 0.858096277 | 0.0879305 | 0.08734 | 0.000591986 | -0.677806514 |
| Rab25 | 0.332786784 | 0.1389058 | 0.13712 | 0.001783779 | -1.300869619 |
| Rad50 | 0.960424116 | 0.4113952 | 0.41097 | 0.000428285 | -0.104213934 |
| Rad51 | 0.024369829 | 0.1550382 | 0.13781 | 0.017233102 | -12.50541433 |
| Raptor | 0.79377184 | 0.762859 | 0.75543 | 0.007433942 | -0.984074094 |
| Rb | 0.533730146 | 0.2241153 | 0.2198 | 0.004313551 | -1.962473631 |
| Rb_pS807_S811 | 0.457540895 | 1.1841248 | 1.11809 | 0.066029881 | -5.90557026 |
| RBM15 | 0.957138438 | 0.4360402 | 0.43545 | 0.000592611 | -0.13609247 |
| Rictor | 0.689051732 | 0.6479663 | 0.6408 | 0.007167023 | -1.11845059 |
| Rictor_pT1135 | 0.017269911 | 0.6853876 | 0.62371 | 0.061679537 | -9.889167527 |

| Protein | Paired TTEST | DMSO Av | TLT Av | Mean diff DMSO-M4 | % Change |
|---|---|---|---|---|---|
| RSK | 0.147394136 | 0.4682432 | 0.45481 | 0.013431948 | -2.953301246 |
| S6_pS235_S236 | 0.09647995 | 1.2531184 | 1.40306 | -0.149946424 | 10.68706343 |
| S6_pS240_S244 | 0.118730496 | 0.8496313 | 0.94022 | -0.090591207 | 9.635081836 |
| SCD | 0.759737499 | 0.0673423 | 0.06663 | 0.000707615 | -1.061931555 |
| SETD2 | 0.041455038 | 0.1748269 | 0.16728 | 0.007542015 | -4.508486479 |
| SF2 | 0.082190928 | 0.0562108 | 0.05337 | 0.002836706 | -5.314761793 |
| Shc_pY317 | 0.413441153 | 0.3421638 | 0.33226 | 0.009902927 | -2.980467043 |
| Smac | 0.20574698 | 0.4343177 | 0.36964 | 0.064677673 | -17.4974742 |
| Smad1 | 0.109784621 | 0.2190075 | 0.20699 | 0.012015241 | -5.804681259 |
| Smad3 | 0.042150885 | 0.5352521 | 0.5145 | 0.020756397 | -4.034318757 |
| Smad4 | 0.109781972 | 0.1975331 | 0.18686 | 0.01066866 | -5.709304748 |
| Snail | 0.021853524 | 0.2435867 | 0.32022 | -0.076637945 | 23.93255874 |
| Src | 0.921787018 | 0.2637837 | 0.26281 | 0.000976124 | -0.371421462 |
| Src_pY416 | 0.208694232 | 0.1672131 | 0.15217 | 0.015041977 | -9.884909758 |
| Src_pY527 | 0.10778136 | 0.700054 | 0.642 | 0.05805016 | -9.042026273 |
| Stat3_pY705 | 0.061361639 | 0.288247 | 0.23968 | 0.048567263 | -20.26340212 |
| Stat5a | 0.004748425 | 0.5811547 | 0.52104 | 0.060117804 | -11.53810864 |
| Stathmin-1 | 0.452980835 | 0.1374042 | 0.14212 | -0.004714313 | 3.317171496 |
| Syk | 0.804047646 | 0.2295709 | 0.22859 | 0.000979491 | -0.428489726 |
| TAZ | 0.293525898 | 0.4116032 | 0.58673 | -0.175121832 | 29.8473411 |
| TFRC | 0.45006093 | 0.5982809 | 0.62051 | -0.022225227 | 3.581790061 |
| TIGAR | 0.203513955 | 0.4447832 | 0.43667 | 0.008111844 | -1.857654354 |
| Transglutaminase | 0.347628428 | 0.3099899 | 0.31823 | -0.008237679 | 2.588612494 |
| TSC1 | 0.724792685 | 0.6995556 | 0.70621 | -0.006652592 | 0.942015755 |
| TTF1 | 0.072765875 | 0.4216622 | 0.40529 | 0.016367572 | -4.038438288 |
| Tuberin | 0.007348867 | 1.2912628 | 1.17141 | 0.119849468 | -10.23118546 |
| Tuberin_pT1462 | 0.472275507 | 0.3208728 | 0.30659 | 0.014281069 | -4.658009245 |
| TWIST | 0.026724298 | 0.0787599 | 0.07013 | 0.00863235 | -12.30950185 |
| Tyro3 | 0.148506776 | 0.2903453 | 0.26374 | 0.026606514 | -10.08820479 |
| UBAC1 | 0.084205337 | 0.4482809 | 0.42428 | 0.023999931 | -5.656613047 |
| UGT1A | 0.129590932 | 0.0798728 | 0.07732 | 0.002550457 | -3.298473152 |
| UQCRC2 | 0.968988613 | 0.097701 | 0.09758 | 0.000124445 | -0.127535891 |
| VEGFR-2 | 0.399973553 | 0.4634747 | 0.45786 | 0.005619443 | -1.227340541 |
| XRCC1 | 0.599402963 | 0.4539372 | 0.45771 | -0.003775075 | 0.824770285 |
| YAP | 0.389819715 | 0.3792817 | 0.38599 | -0.006706443 | 1.737473896 |
| YAP_pS127 | 0.786380304 | 0.7320844 | 0.73928 | -0.007192785 | 0.972948256 |
| YB1 | 0.055196453 | 0.6676773 | 0.7363 | -0.068626495 | 9.320405293 |
| YB1_pS102 | 0.019015082 | 0.6649866 | 0.78032 | -0.11532871 | 14.77975748 |
