## Supplementary Table 5 for "Topsentinol L Trisulfate, a new Marine Natural Product, that Targets Basal-like and Claudin-low Breast Cancers"

| | Topsentinol L Trisulfate EC50 (μM) | | | |
|---|---|---|---|---|
| **Cell Line** | **GFP** | **AMPK** | **CHK1** | **AMPK+CHK1** |
| **HCC1143** | 147.37 | 73.35 | 46.60 | 82.49 |
| **MDA-MB-436** | 88.80 | 81.74 | 61.86 | 57.34 |
